# Diverse Genetic Contexts of HicA Toxin Domains Propose a Role in Anti-Phage Defense

**DOI:** 10.1101/2023.12.08.570793

**Authors:** Kenn Gerdes

**Author notes:** **Address Correspondence:** Kenn Gerdes.

## Abstract

Toxin – antitoxin (TA) modules are prevalent in prokaryotic genomes, often in substantial numbers. For instance, the *Mycobacterium tuberculosis* genome alone harbors close to 100 TA modules, half of which belong to a singular type. Traditionally ascribed multiple biological roles, recent insights challenge these notions and instead indicate a predominant function in phage defense. TAs are often located within Defense Islands, genomic regions that encode various defense systems. The analysis of genes within Defense Islands have unveiled a wide array of systems, including TAs that serve in anti-phage defence. Prokaryotic cells are equipped with anti-phage Viperins that, analogous to their mammalian counterparts, inhibit viral RNA transcription. Additionally, bacterial Structural Maintenance of Chromosome (SMC) proteins combat plasmid intrusion by recognizing foreign DNA signatures. This study undertakes a comprehensive bioinformatics analysis of genetic elements encoding the HicA double-stranded RNA-binding domain, complemented by protein structure modeling. The HicA toxin domains are found in at least 14 distinct contexts and thus exhibit a remarkable genetic diversity. Traditional bicistronic TA operons represent eight of these contexts, while four are characterized by monocistronic operons encoding fused HicA domains. Two contexts involve *hicA* adjacent to genes that encode bacterial Viperins. Notably, genes encoding RelE toxins are also adjacent to Viperin genes in some instances. This configuration hints at a synergistic enhancement of Viperin-mediated anti-phage action by HicA and RelE toxins. The discovery of a HicA domain merged with an SMC domain is compelling, prompting further investigation into its potential roles.

**Importance:** Prokaryotic organisms harbor a multitude of Toxin – Antitoxin (TA) systems, which have long puzzled scientists as “genes in search for a function”. Recent scientific advancement have shed light on a primary role of TAs as anti-phage defense mechanisms. To gain an overview of TAs it is important to analyze their genetic contexts that can give hints on function and guide future experimental inquiries. This manuscript describes a thorough bioinformatics examination of genes encoding the HicA toxin domain, revealing its presence in no fewer than 14 unique genetic arrangements. Some configurations notably align with anti-phage activities, underscoring potential roles in microbial immunity. These insights robustly reinforce the hypothesis that HicA toxins are integral components of the prokaryotic anti-phage defense repertoire. The elucidation of these genetic contexts not only advances our understanding of TAs but also contributes to a paradigm shift in how we perceive their functionality within the microbial world.

## Introduction

Prokaryotes and their mobile genetic elements, such as phages and plasmids, have been locked in a co-evolutionary arms race spanning billions of years. In this intricate interplay, bacteria and archaea have developed an arsenal of innate and acquired defenses against phages and plasmids. Acquired immunity, typified by CRISPR-Cas systems, relies on the memory of prior phage encounters. In contrast, innate defenses, like restriction-modification systems, are hardwired to indiscriminately degrade invasive genetic material. The concept of "Defense Islands" has been pivotal in recent discoveries by uncovering genomic regions in which defense genes are clustered (<u>1-3</u>). These revelations have unveiled a diverse array of defense strategies, demonstrating evolutionary links between prokaryotic and mammalian innate immunity systems. For instance, Theoresis phage defense systems possess domains akin to Toll-like receptors, and prokaryotic Viperins (pVips), like their mammalian orthologues, inhibit phage transcription by generating modified nucleotides (<u>4</u>). Similarly, anti-plasmid mechanisms employ SMC ATPases, which recognize signatures in foreign plasmid DNA and prevent plasmid establishment (<u>1</u>, <u>5</u>) (<u>Robins WP, Mekalanos JJ et al. BioRxiv 2023</u>). Toxin – antitoxin (TA) modules, often situated in Defense Islands (<u>1-3</u>, <u>6</u>), have only recently been associated with anti-phage activity, particularly via an abortive infection mechanism which halts infection at the cost of the host cell, thereby protecting the clonal population (<u>7-9</u>)(<u>Smith, Foster et al. NRM 2023, in press</u>). Recent investigations strongly support the notion that TAs commonly function as anti-phage elements (<u>10-23</u>).

TA modules are categorized into different Types based on how the antitoxin counteracts the toxin. In Type I and III systems, small RNAs act as antitoxins by either blocking toxin translation or binding directly to the toxin, while Type II systems use protein antitoxins to achieve neutralization by direct protein-protein contact (<u>24</u>, <u>25</u>). Based on toxin sequence similarity, the different Types of TAs have been subdivided into families. For example, Type II TAs encode RelE, MazF, VapC and HicA family toxins. All these toxins are RNases (also called mRNA interferases) that inhibit translation by cleavage of mRNA, rRNA or tRNA and may thus induce abortive infection upon activation.

HicA toxins constitute a large family of small, mono-domain RNases ranging 50 to 100 amino acids. They are found in both bacterial and archaeal species, often in multiple copies per genome (<u>26</u>, <u>27</u>). The core of HicA RNases exhibits the characteristic α-β-β-β-α topology of the double-strand RNA Binding Domain (dsRBD) fold (**Figure 1A**) (<u>27-32</u>). Superimposition of the crystal structures of *Escherichia coli* K-12 HicA and HicA of *Thermus thermophilus* HB8 known to exhibit the dsRBD topology (<u>27</u>) exposes their highly related tertiary structures (**Figure 1B**).

**Figure 1.**
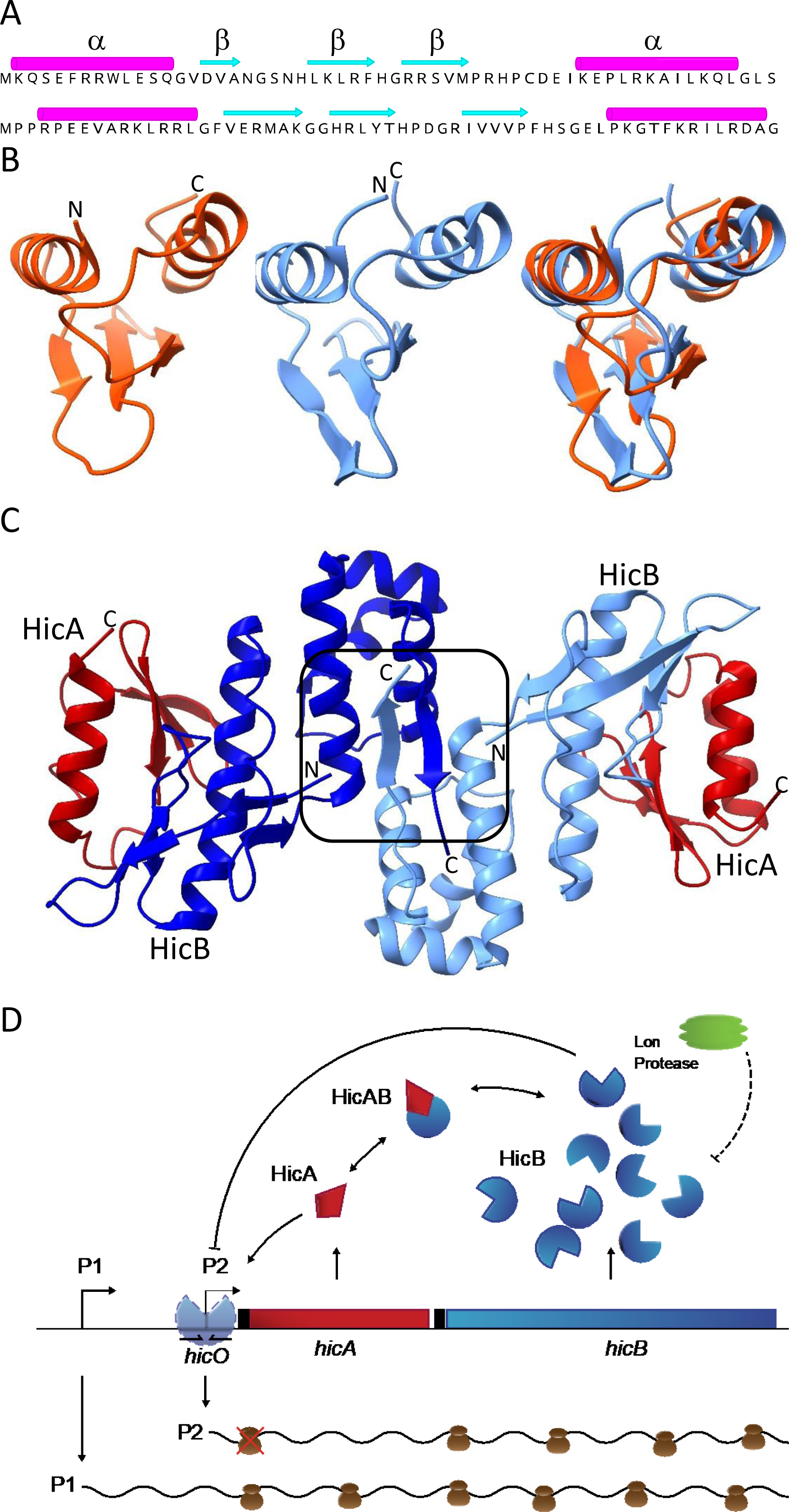
Components and regulatory features of the *E. coli hicAB* toxin – antitoxin module. (**A**) Secondary structures of HicA of *E. coli* K-12 (upper) and HicA of *T. thermophilus* TTHA1913. (**B**) Tertiary structures of *E. coli* K-12 HicA (left, orange), *T. thermophilus* TTHA1913 HicA (middle, light blue) and their superimposition (right). In the superimposition of the two HicA structures, the RMSD between 36 pruned atom pairs is 1.3Å. The *E. coli* HicA structure was determined experimentally (<u>31</u>) while that of *T. thermophilus* was modelled by AlphaFold2. (**C**) Tertiary structure of the *E. coli* K-12 heterotetrameric HicA_2_B_2_ complex determined experimentally. The two HicAs are shown in red while the two HicBs are shown in blue and light-blue. The HicB HTH DNA-binding motifs that dimerizes HicB are boxed-in. The **Figure** was adapted from (<u>31</u>). (**D**) Proposed regulation of the *E. coli hicAB* operon (<u>33</u>). The *hicAB* operon is transcribed by two promoters, P1 and P2. During steady-state growth-conditions, P1 is constitutive while P2 is repressed by HicB binding to the *hicO* operator. Excess HicA toxin ([HicA] > [HicB]) destabilizes HicBs binding to *hicA* and excess of HicA therefore leads to activation of P2. Notably, the *hicAB* transcript synthesized by P2 produces HicB but not HicA. Therefore, the HicA-mediated derepression of *hicAB* transcription specifically stimulates synthesis of HicB but not HicA under conditions of excess HicA. The *hicAB* operon of *Burkholderia pseudomallei* is regulated by a related mechanism: excess of HicA destabilizes the binding of HicB to the operator in the promoter region (<u>32</u>). Symbols: Arrows indicate stimulation, and lines ending in a bar symbolize inhibition or protein degradation (Lon hexamer shown in green degrades HicB (<u>26</u>) shown in blue; HicA is shown in red). *hicO* symbolizes the inverted repeat to which HicB dimers bind and repress transcription. Messenger RNAs are shown as wavy lines and ribosomes as brown bodies. A red cross-over symbolizes that the *hicA* cistron of the P2-generated transcript is not translated.

The HicA toxins that have been examined experimentally are all encoded by bicistronic *hicAB* operons where the downstream gene encodes a HicB antitoxin. In these systems, HicB comprises an N-terminal partial RNase H domain and a C-terminal DNA-binding-domain (DBD) of the HTH or the RHH type (<u>27</u>, <u>31</u>, <u>32</u>). The partial RNase H domain of HicB exhibits a β-β-β-α-β-α topology, with the first four secondary structure elements characteristic of partial RNase H folds (**Figure S1A**) (<u>27</u>). The superimposition of *E. coli* K-12 HicB’s partial RNase H fold on that of TTHA1013 from *T. thermophilus* HB8, known to possess a partial RNase H fold, confirms the similarity of HicB’s fold (**Figure S1B**).

Crystallographically, HicA and HicB of *E. coli* K-12 form an A_2_B_2_ heterotetrameric complex (<u>31</u>) (**Figure 1C**). HicB interacts with HicA by packing of helix α1 of the partial RNase H motif against the β sheet of HicA (**Figure 1C**), mirroring the observed behavior in RHH-containing HicAB complexes *of Burkholderia pseudomallei* and *Streptococcus pneumoniae* (<u>30</u>, <u>32</u>). This implies that HicA inhibition by HicB operates independently of the type of HicB DNA-binding domain. HicB binds to palindromic operators in the *hicAB* operon promoter region via its C-terminal DBD, thereby autorepressing transcription (**Figure 1D**). Notably, high levels of HicA destabilize the HicAB-DNA complex and thereby stimulates operon transcription (<u>32</u>, <u>33</u>). This phenomenon, observed in many other type II TA families, underscores the intricate repression and derepression mechanisms of *hicAB* operons (<u>34-37</u>). Further details of *hicAB* operon regulation and derepression are described and visualized in **Figure 1D**.

Motivated by the exponential growth of microbial DNA databases, this study undertakes a comprehensive bioinformatics analysis of genetic elements encoding HicA dsRBD domains. The findings reveal the presence of HicA domains in at least 14 distinct genetic contexts, eight of which adhere to the canonical bicistronic TA operon configuration. Remarkably, only two of these genetic contexts have undergone experimental analysis. Four configurations encompass monocistronic operons, featuring fused HicAB domains, while the remaining two configurations involve *hicA* genes in operon with bacterial Viperins, which serve as guardians against bacteriophage invasions (<u>4</u>). The most common bicistronic operon structure is *hicBA* in which antitoxin HicB does not have a DBD, raising the question of how transcription of these operons is regulated. Lastly, the discovery of a HicA domain fused to a Structural Maintenance of Chromosome (SMC) domain raises intriguing functional questions.

## Results and Discussion

### Fourteen Classes of HicA Domains

Through database searches utilizing experimentally verified HicA toxin sequences, a striking diversity of genetic contexts encoding HicA domains emerged. Automated inspection of the neighboring sequences led to the classification of HicA domains into 14 distinct sequence classes (**Figure 2**; **Table S1**). Each automated gene annotation was meticulously validated by manual examination of the DNA sequences. The 14 classes encompass a wide spectrum of genetic organizations, with Classes 1 to 10 featuring bicistronic operons, resembling the genetic arrangement of typical bicistronic TA modules. In contrast, Classes 11 to 13 consist of monogenic operons that encode fused HicA and HicB domains, while Class 14 is an interesting case of an SMC domain fused to a HicA domain. Notably, HicA-encoding genes exhibit a ubiquitous presence across prokaryotic phyla, underscoring their prevalence in the prokaryotic realm (**Table S1**). However, it is noteworthy that Classes 3 and 4 are relatively scarce in Archaea, and archaeal HicA-encoding TA loci often belong to Class 1 or 2. The following sections describe the distinctive features of these diverse classes.

**Figure 2.**
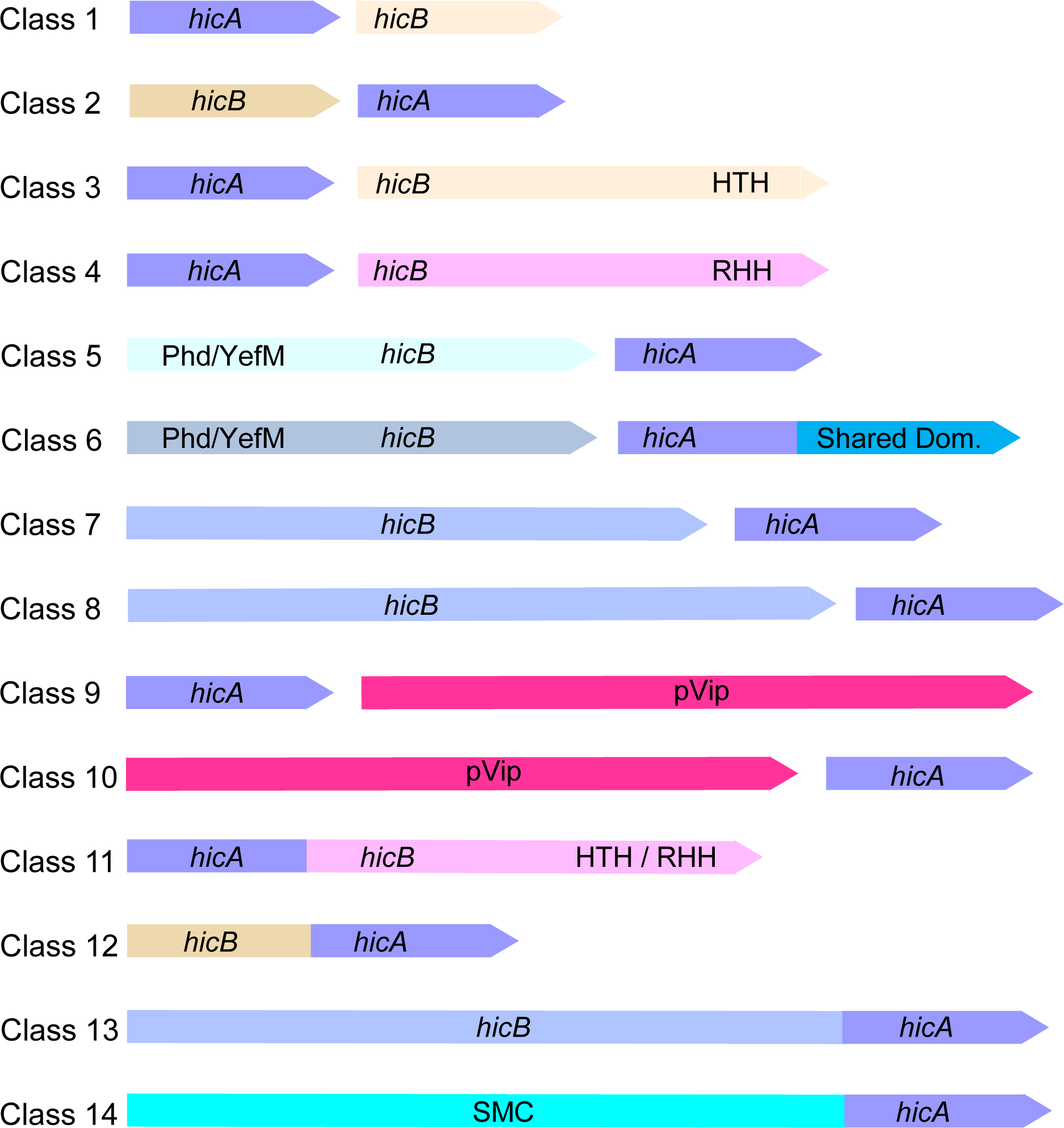
Fourteen distinct genetic contexts encoding HicA domains. Genetic organization of 14 HicA-domain classes derived from data in **Table S1** (Sheets 1 to 14). **Class 1:** *hicAB* in which HicB is small (67 – 97 a a) and devoid of a recognizable DBD; **Class 2:** *hicBA* with a reversed gene order compared to that of Class 1 in which *hicB* also is small (63 – 103 codons) and devoid of a recognizable DBD; **Class 3:** *hicAB* in which HicB has a C-terminal HTH domain; **Class 4:** *hicAB* in which HicB has a C-terminal RHH domain; **Class 5:** *hicBA* with a reversed gene order compared to Class 3 and 4 and in which *hicB* has an N-terminal Phd/YefM DBD; **Class 6:** Similar to Class 5 but HicA has an extended C-terminal domain called the Shared Domain; **Class 7:** *hicBA* loci in which *hicB* is larger than *hicB* of the previous classes (216 – 233 codons); **Class 8:** *hicBA* loci in which *hicB* is even larger than *hicB* of Class 7 (331 – 356 codons); **Class 9:** *hicA* upstream of a gene encoding a pVip; **Class 10:** *hicA* downstream of a gene encoding a prokaryotic Viperin (pVip); **Class 11:** *hicAB* monocistronic operon encoding a HicA domain fused to HicB with a C-terminal DBD (HTH or RHH); **Class 12:** *hicBA* monocistronic operon encoding a HicA domain fused to a small HicB domain; **Class 13:** *hicBA* monocistronic operon encoding a HicA domain fused to a large HicB domain; **Class 14:** A Structural Maintenance of Chromosome (SMC) domain fused to a C-terminal HicA domain.

### Class 1 and 2: Small and Compact *hicAB* Modules

Classes 1 and 2 comprise *hicAB* and *hicBA* operons, respectively, with HicBs encoded by these modules characterized by their diminutive size (ranging from 57 to 94 amino acids) and the absence of an identifiable DNA binding domain (DBD) (**Figure 2**). Consequently, these TA modules are exceptionally compact, typically encoding HicAs of 60 to 85 amino acids and HicBs of 60 to 90 amino acids. In most instances, the *hicA* and *hicB* genes of both Class 1 and Class 2 are closely linked or exhibit overlap, suggesting translational coupling (**Table S1**). The lack of DBDs of HicB is significant because it raises questions regarding the regulatory mechanisms governing the expression of the operons. It can be argued that Class 1 HicBs, devoid of a DBD, are non-functional genes arising from premature stop-codon mutations (**Figure 2**). While database searches indeed uncovered instances of such cases, a thorough sorting of Class 1 HicBs using BLASTP analyses supports the functionality of most Class 1 *hicAB* modules, as explained in detail in the Materials and Methods section.

Notably, toxins and antitoxins of Classes 1 and 2 have not yet undergone experimental analysis. Structure modeling unveiled that Class 1 and Class 2 HicA toxins from a cyanobacterial phage and *Klebsiella pneumoniae* feature the canonical α-β-β-β-α fold (**Figure 3A**, **3C**), as first reported for the HicA double-strand RNA binding domain (dsRBD) (<u>27</u>). The corresponding HicA•HicB dimer structures are shown in **Figure 3B** and **3D**, respectively. Interestingly, phage-encoded HicA in complex with its cognate HicB is predicted to undergo a conformational change, potentially resulting in the loss of its N-terminal β-sheet (**Figure 3B**). In contrast, HicA of *K. pneumoniae* does not exhibit such a change in configuration in the predicted complex (**Figure 3D**). One possibility is that in the phage HicAB complex, HicA changes configuration and thereby loses its RNase activity – that is – HicA’s interaction with HicB leads to a structural change that inactivates the enzyme. Alternatively, the predicted structural change might be an artifact of the modeling process, necessitating further investigation to elucidate this issue. Importantly, both HicBs of the HicA•HicB complexes exhibit the canonical β-β-β-α-β configuration of the partial RNase H fold (<u>27</u>) (**Figure 3B**, **3D**). The AlphaFold2 modeling produced high-quality structures, further confirmed by ModFOLDdock analysis, yielding high Assembly and Interface quality scores (**Figure 3E**, **3F**).

**Figure 3.**
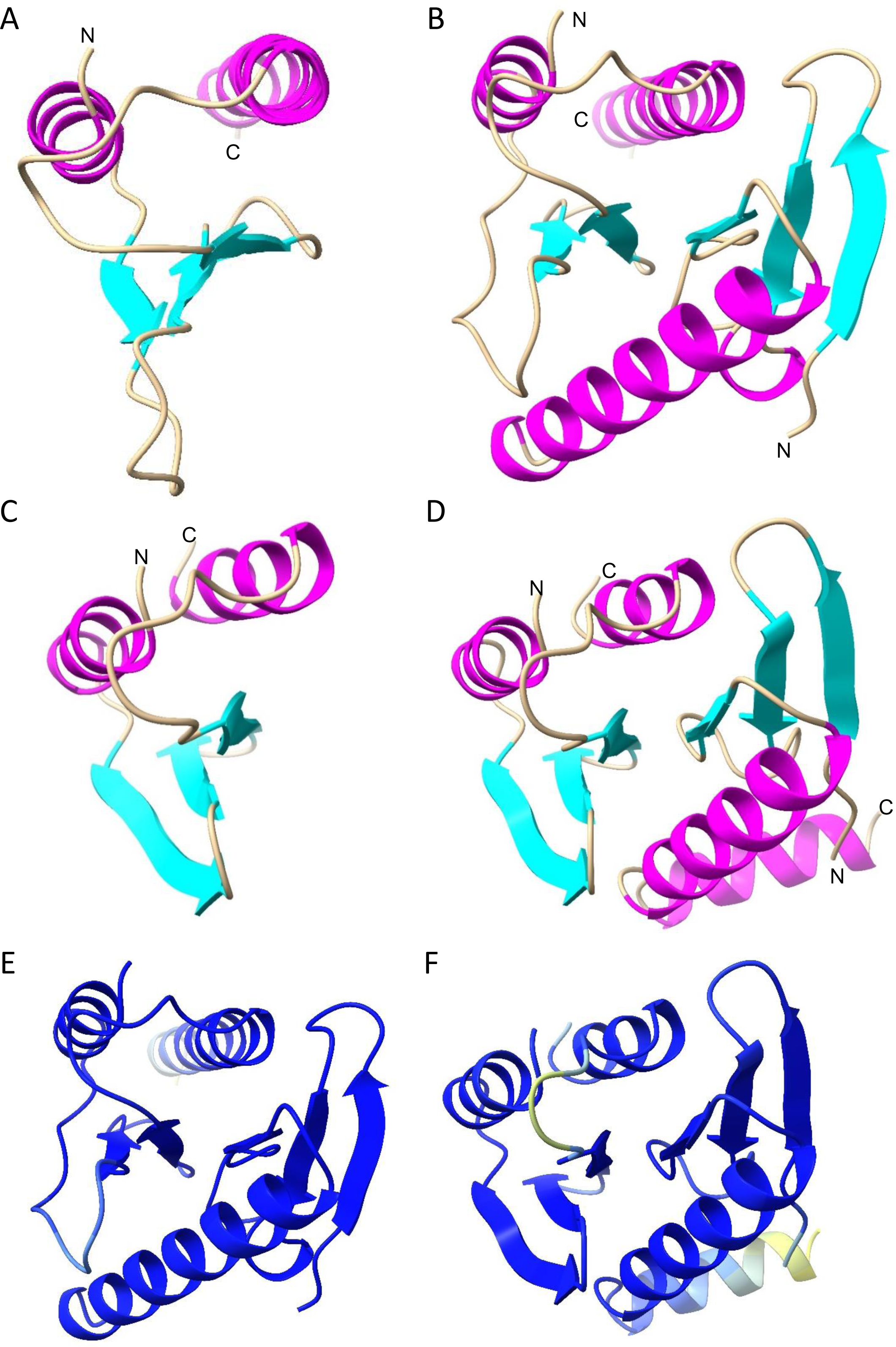
Modelling of Class 1 and 2 HicA monomer and HicAB dimer structures. (**A**) Structure model of Class 1 HicA monomer of *Planktothrix* phage PaV-LD (YP_004957304.1) generated by AlphaFold2. (**B**) Structure model of Class 1 HicAB dimer of *Planktothrix* phage PaV-LD (YP_004957304.1, YP_004957303.1) generated by MultiFOLD. Dimer plDDT: 0.969; pTM: 0.903.; Assembly quality: 0.9503; Interface quality: 0.9524. Assembly and Interface quality were calculated by ModFPOLDdock. (**C**) Structure model of Class 2 HicA monomer of *Campylobacter* sp. RM12654 (MBZ7977437.1) generated by AlphaFold2. (**D**) Structure model of Class 2 HicAB dimer of *Campylobacter* sp. RM12654 (MBZ7977437.1, MBZ7977438.1) generated by MultiFOLD. Dimer plDDT: 0.925; pTM: 0.850.; Assembly quality: 0.9125; Interface quality: 0.9069. Assembly and Interface quality were calculated by ModFOLDdock. (**E**) and (**F**) Dimer structure models from (B) and (D) colored according to AlphaFold2 quality scheme (blue represents high quality, yellow low quality).

### Class 3 and 4: Classical *hicAB* Operons with HTH or RHH DNA-Binding Domains

Classes 3 and 4 encompass the model *hicAB* operons, with Class 3 encoding HicB antitoxins with a C-terminal helix-turn-helix (HTH) DBD and Class 4 HicB having a C-terminal ribbon-helix-helix (RHH) DBD (**Figure 2**). As elaborated in the Materials and Methods section, all DNA binding domains were rigorously validated using AlphaFold2, FoldSeek, Phyre2, or, in a few ambiguous cases, by sequence similarity searches (BLASTP). In experimentally analyzed modules, HicBs employ their HTH or RHH domains to autorepress transcription of their cognate *hicAB* operon by binding to palindromic operators in the promoter regions. Simultaneously, the antitoxins counteract the detrimental RNase activity of HicA (as illustrated in **Figure 1D**). The distances between *hicA* and *hicB* in these operons vary: a substantial portion (31% of 119) of Class 3 genes are closely linked (with a separation of ≥ 10 bases between *hicA* and *hicB*; **Table S1**), and a similar trend is observed in Class 4 genes (41% of 171). Several experimental structures of the components encoded by Class 3 and 4 operons have been elucidated and will not be explored further here (<u>28-32</u>).

### Class 5 and 6: HicBA Modules with Unconventional Arrangements

Classes 5 and 6 represent a distinctive departure from the typical *hicBA* genetic organization. These classes feature a reversed gene order compared to the classical Class 3 and 4 *hicBA* modules and encode HicB antitoxins with a Phd/YefM DNA-binding domain (DBD) in their N-termini (**Figure 2 and 4A**). Similar to the HTH and RHH domains of HicBs of Classes 3 and 4, the Phd/YefM DBD of Classes 5 and 6 HicBs dimerizes the complex and likely function to regulate transcription of their cognate *hicBA* operons via binding to operators in the promoter regions (**Figure 4B**, **4C**). Most *hicBA* genes belonging to Classes 5 and 6 are closely linked and in many cases overlap, indicating translational coupling. Classes 5 and 6 introduce a novel feature of *hicBA* operon regulation by utilizing Phd/YefM DBDs.

**Figure 4.**
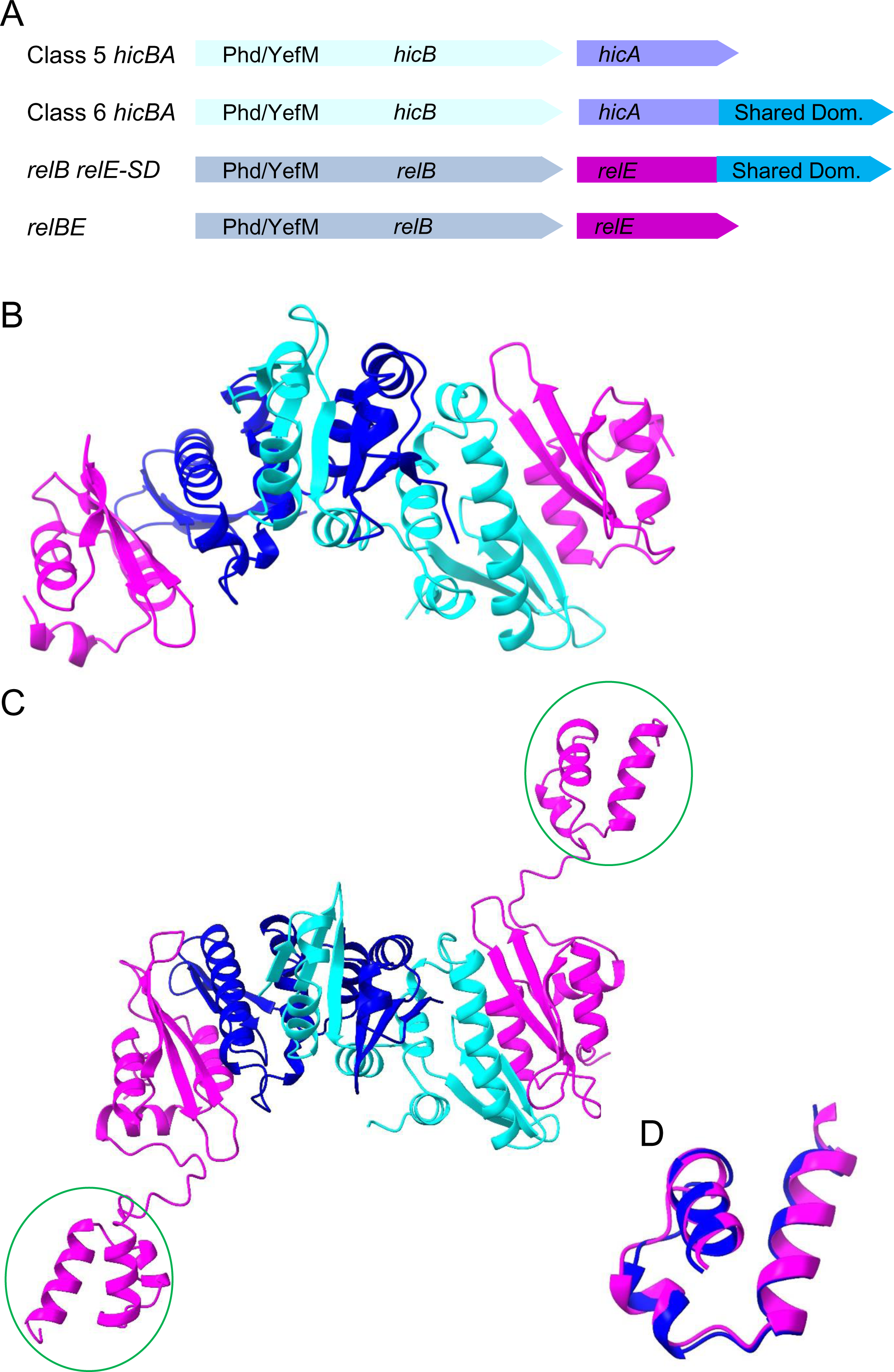
Genetic organizations and structures of with Shared Domains. (**A**) Comparison of the genetic organizations of HicA and RelE toxins with Shared Domains. Class 5 and 6 *hicA* are in operons with a downstream *hicB* encoding Phd/YefM DBD domains. RelE with Shared Domains also have cognate antitoxins with Phd/YefM DBD encoded by downstream *relB* genes. *relB* genes encoding Phd/YefM DBD in their N-terminal parts were identified using webFlaGs. (**B**) Structure of tetrameric Class 5 HicA_2_B_2_ complex (WP_078438722.1, WP_217700609.1). HicB dimerizes via the N-terminal Phd/YefM DBD and interacts with HicA monomers mainly via its central β-sheet of the partial RNase H-fold that aligns with α-helix 2 of HicA. (**C**) Structure of tetrameric Class 6 HicA_2_B_2_ complex (MBB5067807.1, MBB5067806.1). Again, HicB dimerizes via the N-terminal Phd/YefM DBD and interacts with HicA monomers mainly via its central β-sheet that aligns with α-helix 2 of HicA. The Shared Domains of the two HicA monomers are indicated with a green circle. (**D**) Superposition of Shared Domains of Class 6 HicA (MBB5067807.1) and a RelE toxin (WP_141355914.1) with a Shared Domain (From **Figure S4**). The superposition has an RMSD of 0.838Å across the entire structures (38 pairs).

HicAs from Classes 1 to 5 are small, mono-domain proteins featuring a dsRNA-binding domain (dsRBD) fold. However, Class 6 HicAs have an additional domain, approximately 55 to 60 amino acids in size, situated at their C-termini, referred to here as the "Shared Domain" (**Figure S2**). Notably, conserved prolines found at the junction between the N-terminal HicA dsRBD domain and the Shared Domain may act as "domain-breakers" maintaining separation between the N and C-terminal domains (**Figure S2**). While Class 6 HicAs are predominantly present in Actinomycetes, they also occur in other phyla.

BLAST analyses led to the identification RelE/ParE toxins also consisting of two domains where the C-terminal domain exhibits sequence similarity to the Shared Domain of Class 6 HicAs. Sequence alignments of RelE/ParE toxins with and without a Shared Domain reveals a pattern very similar to that of the alignments of Class 5 and 6 HicAs (**Figure S3**). Alignment of the Shared Domains of Class 6 HicAs and RelE/ParE toxins demonstrate their sequence similarity and their conserved secondary structure (**Figure S4**). The genetic organization of Class 6 *hicBA* and *relBE* with a Shared Domain are strikingly similar, suggesting a common function of the Shared Domains of the two toxin families (**Figure 4A**).

The structural modelling of the tetrameric Class 5 HicA_2_B_2_ complex show that HicB dimerizes through the Phd/YefM domains of HicB by domain-swapping (**Figure 4B**). The two HicAs interact solely via each HicB subunit and do not interact themselves. Hence, the Class 5 HicA_2_B_2_ complex exhibits a compact structure that in this respect resembles the crystal structure of the HicA_2_B_2_ complex of *E. coli* K-12 (<u>31</u>) (**Figure 1C**). In the Class 6 HicA_2_B_2_ complex, the modelled structure remains largely similar, with the exception of the Shared Domain connected to the dsRBD of HicA through a long, flexible linker (**Figure 4C**). Notably, the Shared Domain of Class 6 HicAs, consisting of three α-helices, extends outward from the compact structure and may thus be available for interactions with external factors.

A similar theme arises with RelB_2_E_2_ complexes in which RelE is extended by a C-terminal Shared Domain. As seen from **Figure S5A**, the canonical RelB_2_E_2_ complex of *E. coli* K-12 exhibits a V-shaped structure generated by dimerization via the RHH domains of RelB and RelB domain-swapping (<u>38</u>). The RelE subunits interact with the C-terminal domain of RelB. A RelB_2_E_2_ complex in which RelE has a C-terminal Shared Domain complex in which two RelBs dimerize by domain-swapping via their Phd/YefM domains forming a structure in which the two RelEs interact with the C-terminal parts of RelB (**Figure S5B**). Again, the Shared Domain, consisting of three α-helices, extends outward from the complex suggesting potential interactions with external factors. Superposition of the Shared Domains from a RelE and a Class 6 HicA reveals a remarkable root mean square deviation (RMSD) of 0.838Å, indicating a common ancestral origin. (**Figure 4D**). The presence of domains shared across distinct toxin families is unique and deserves experimental scrutiny.

### Class 7 and 8: Remarkable Diversity of *hicBA* Modules

These classes are characterized by encoding relatively long HicB antitoxins, with Class 7 HicBs spanning 216 to 233 amino acids, and Class 8 HicBs extending to 342 to 356 amino acids (**Figure 2**). In line with typical TA modules, the *hicA* and *hicB* genes within these classes are closely linked, and in many instances overlap (**Table S1**). For Class 7, structural modeling was employed to gain insight into the HicA and HicB components. The Class 7 HicA, encompassing a typical dsRBD, signifies its potential role as an RNase (**Figure S6A**). On the other hand, Class 7 HicB comprises two domains separated by an α-helix (**Figure S6B**). FoldSeek searches identified the presence of DUF1902 domains in both the N and C-terminal parts of HicB. DUF1902 domains are characterized by the presence of an α-helix and four β-strands. (**Figure S6B**). Using webFlaGs, it became apparent that many DUF1902-encoding genes are juxtaposed to a *hicA* gene. This insight provides a potential avenue for further investigation into the significance of DUF1902 domains. A Class 7 HicAB dimer model further supported the notion that HicA likely interacts with the antiparallel β-sheets of the C-terminal domain of HicB (**Figure S6C**).

Class 8 introduces additional complexity, with a longer Class 8 HicA of ca. 95 amino acids when compared to canonical mono-domain HicA toxins (Classes 1 to 5). Class 8 HicA retains the characteristic α-β-β-β-α dsRBD fold but features two additional small α-helices at the C-terminus (**Figure S7A**). Importantly, secondary structure predictions based on a sequence alignment of Class 8 HicAs revealed the conservation of these two α-helices (**Figure S7C**).

Class 8 HicB exhibits even greater complexity, characterized by three distinct domains (**Figure S7B**). The middle domain (aa 175 to 253) shares structural similarities with lysyl-tRNA synthetases, the C-terminal domain (aa 255 to 342) displays the HicB-fold, and the N-terminal domain (aa 1 to 171) does not exhibit significant similarity to domains with known functions. Alignment of HicB sequences confirmed the three-domain structure (**Figure S8**). The structural similarities found in Class 8 HicB domains opens avenues for further exploration, especially regarding the functional role of the middle domain that resembles lysyl-tRNA synthetases. Notably, neither HicA nor HicB of Classes 7 and 8 feature a recognizable DNA-binding domain (DBD), again raising questions of how synthesis of these proteins is regulated.

### Class 9 and 10: *hicA* and *relE* are Adjacent to Prokaryotic Viperin Genes

Eukaryotic Viperins are antiviral proteins that modify CTP and thereby cause termination of viral RNA synthesis (<u>39</u>, <u>40</u>). In turn, infection by a broad range of RNA and DNA viruses is inhibited. Prokaryotic Viperins are orthologues of eukaryotic Viperins that have anti-phage activity (<u>4</u>). Notably, pVip antiviral activities extend beyond CTP to include the modification of GTP and UTP, making them powerful defenders against phage infections (<u>4</u>).

As also noted by others (<u>4</u>), *hicA* genes are found adjacent to genes encoding pVIPs, presenting a fascinating convergence (**Figure 2**). Genes encoding HicAs are located both upstream (Class 9) and downstream (Class 10) of pVIP genes, in both cases adjacent to the pVIP-encoding gene (**Table S1**). The *hicA* genes located upstream and downstream of the pVip genes all exhibit the typical HicA secondary (α-β-β-β-α) and tertiary structures, suggesting that at least some of these HicAs are active toxins (**Figure S9**).

A further fascinating revelation emerged during the exploration of *hicA* genes linked to pVIPs: RelE-encoding genes are also closely associated with pVIP-encoding genes (**Table S1** and **Figure S10A**). The RelEs that accompany pVIPs adhere to the typical RNase I fold characteristic of RelE-homologous RNases (**Figure S10B**, **S10C**). This finding underscores the possibility that both HicA and RelE RNases operate in tandem with pVIPs to mount a robust defense against phage infections.

When activated, canonical HicA and RelE RNases arrest translation (<u>26</u>, <u>41</u>, <u>42</u>). In many cases, inhibition of translation by TA-encoded toxins contributes significantly to phage defense by facilitating abortive infection, a phenomenon observed with both Type 2 and Type 3 TA systems (<u>9</u>, <u>19</u>, <u>43</u>, <u>44</u>) (and additional references op. cit.). The genetic linkage of pVips and two evolutionary independent RNase families raises several important questions, in particular, how is production and activity of the toxic RNases regulated and do the RNases contribute to the antiviral activity of pVips?

### Class 11 to 13: Distinct Features of Fused HicAB and HicBA Toxin-Antitoxins

Classes 11 to 13 have a unique configuration where HicA and HicB domains are fused, forming monocistronic operons, a notable departure from the typical bicistronic structure in other classes (**Figure 2**). Particularly, Class 11 HicBs have DNA-binding domains (HTH or RHH) at the C-termini, reminiscent of the HicBs of Classes 3 and 4 (Table S1).

Molecular modelling shows that Class 11 HicBAs have three domains: a canonical N-terminal HicA domain, a middle RNase H domain, and a C-terminal DNA-binding domain (RHH or HTH) as illustrated in **Figure 5A**, **5C**. Both RHH and HTH domain-containing subclasses form dimers, a critical feature for DNA binding (**Figure 5B, 5D**). The superposition of experimental RHH and HTH structures with Class 11 models, detailed in **Figure S11**, supports this dimerization through DNA-binding domains.

**Figure 5.**
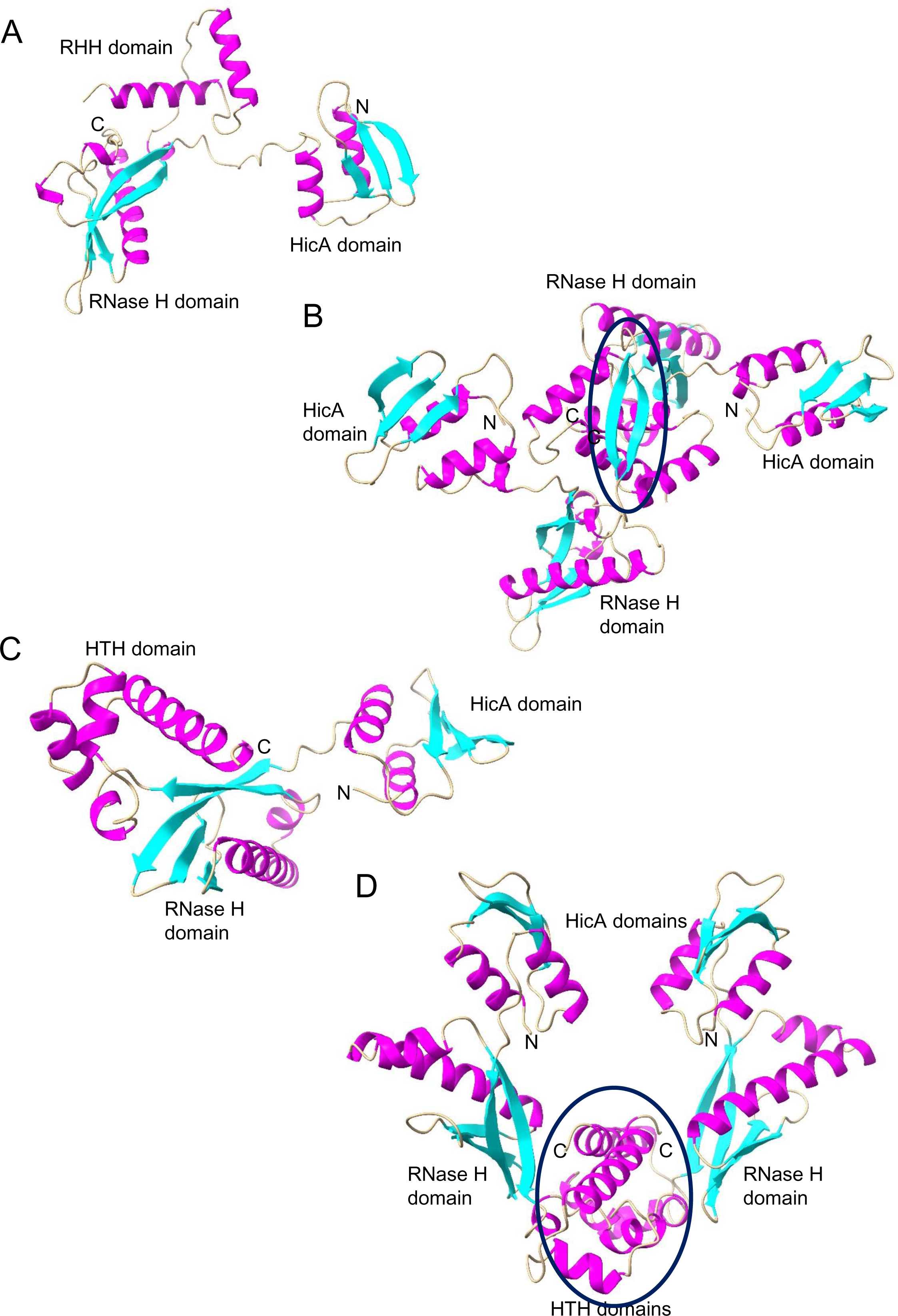
Structure Models of Class 11 fused HicAB mono-domain toxin – antitoxins. (**A**) Monomer model of Class 11 fused HicAB (KPW96986.1) generated by AlphaFold2 with its three domains marked up. (**B**) Dimer model of Class 11 fused HicAB (KPW96986.1). The monomers dimerize via their RHH domains (dark blue ellipse). (**C**) Monomer model of Class 11 fused HicAB (WP_243550782.1). (**D**) Dimer model of Class 11 fused HicAB (WP_243550782.1) model of Class 11 HicAB (WP_243550782.1). The monomers dimerize via their RHH domains (dark blue ellipse).

Fused Class 12 HicBA resemble Class 2 HicBA in terms of gene orientation and protein sizes (**Figure 2**). This structural similarity is reinforced through molecular modeling, which reveals that the HicA domain of a Class 12 HicBA aligns remarkably well with a Class 2 HicA (**Figure 6A**, **6C** and **6E**), with a notable RMSD of only 0.907Å. Similarly, the HicB domains of these classes align closely (RMSD of 0.779Å), hinting at an evolutionary linkage (**Figure 6B, D, F**). This suggests a potential evolutionary trajectory involving the fusion of ancestral separate *hicBA* genes, although the reverse scenario cannot be excluded.

**Figure 6.**
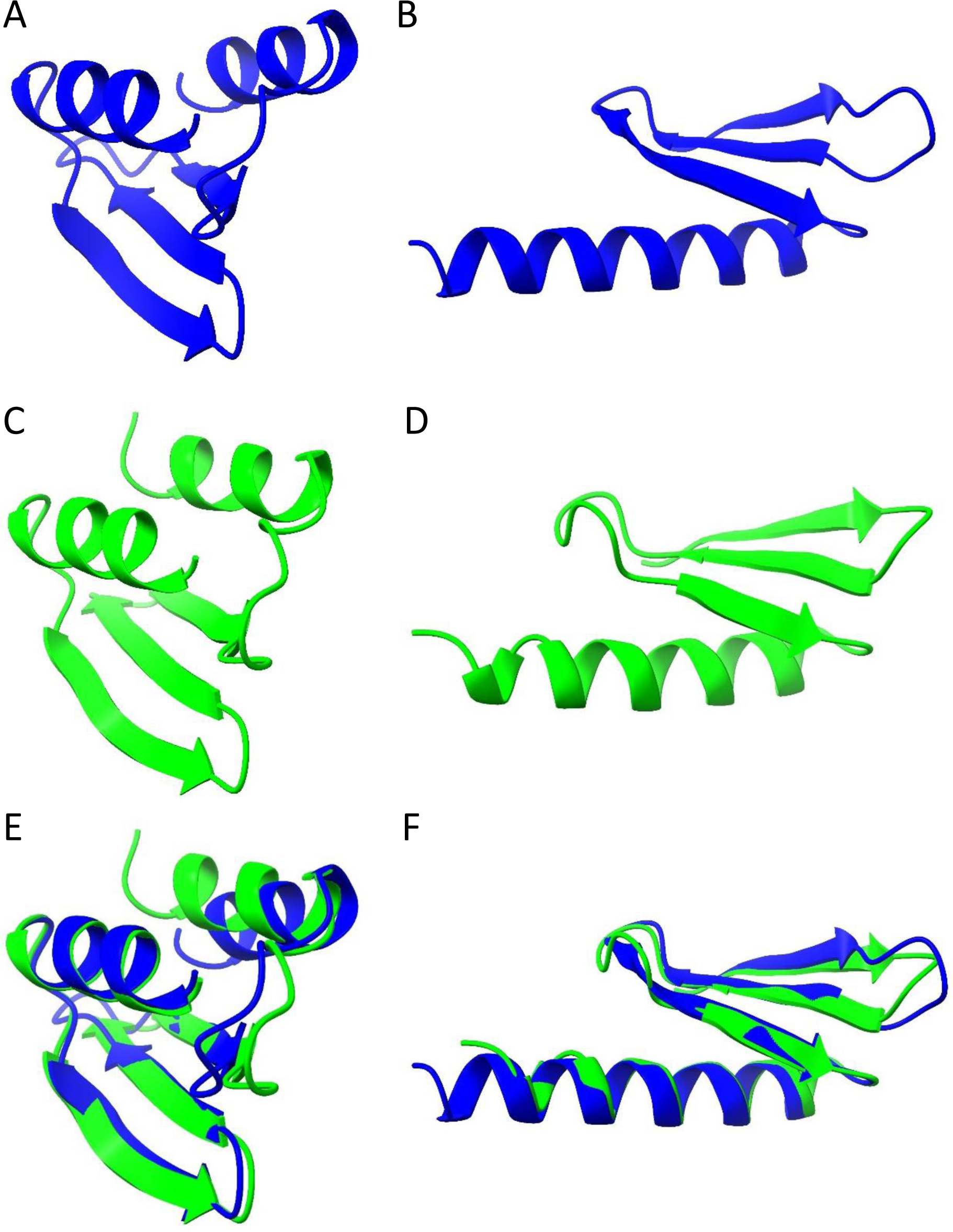
Comparison of Class 2 and 12 HicBA structure models. (**A**) Class 12 HicA domain in blue (aa 74 to 127 of MBI3260764.1; #5 in Sheet 12 of **Table S1**). (**B**) Class 12 HicB domain in blue (aa 12 to 60 of MBI3260764.1; #5 in Sheet 12 of **Table S1**). (**C**) Class 2 HicA in green (aa 9 to 63 of MBZ7977437.1; #129 in Sheet 2 of **Table S1**). (**D**) Class 2 HicB in green (aa 2 to 46 of MBZ7977438.1; #129 of Sheet 2 in **Table S1**). (**E**) Superimposition of structures in (A) and (C) (RMSD between 48 pruned atom pairs is 0.907Å). (**F**) Superimposition of structures in (B) and (D) (RMSD between 46 pruned atom pairs is 0.779Å). All structure models were generated by AlphaFold2 and annotated in ChimeraX.

The functional aspects of Class 12 HicBA was addressed using a phylogenetic analysis comparing artificially fused Class 2 HicBAs with naturally fused Class 12 HicBAs (**Figure S12**). Notably, several Class 12 HicBAs cluster distinctly, indicating that the proteins evolve under selection pressure. This observation indicates that at least some of the Class 12 HicBAs are functional.

Class 13 HicBAs are larger (340 to 440 aa) and feature a unique domain arrangement with predominantly α-helices in the N-terminal region and a C-terminal domain following the canonical HicA configuration (**Figure S13A, S13B**).

HicA belonging to Classes 11 to 13 all share the "fused" configuration and in this respect obviously departs from the canonical Type II TA paradigm. This unusual organization raises the question of how the toxin activity of the fused TAs is regulated. Importantly, in this connection, some fused Type II TAs have been investigated experimentally. The fused TA system CapRel^SJ46^ protects *E. coli* against diverse phages (<u>13</u>). Notably, the C-terminal domain of CapRel^SJ46^ regulates the toxic N-terminal region, serving as both antitoxin and phage infection sensor. After infection by certain phages, newly synthesized major capsid protein of the phage binds directly to the C-terminal domain of CapRel^SJ46^ to relieve autoinhibition, enabling the toxin domain to pyrophosphorylate tRNAs, which blocks translation and thereby restricts viral infection. It is thus possible that the fused HicABs function by similar or related mechanisms to curb phage infection. In addition, the EzeT TA protein of enterobacterial strains consists of two domains in which the C-terminal domain is a toxic kinase while the N-terminal domain inhibits the kinase activity and thereby functions as a cis-acting antitoxin (<u>45</u>).

### Class 14: Novel Fusion of SMC and HicA Domains

Class 14 introduces a surprising fusion of SMC and HicA domains, a combination not previously observed in prokaryotic proteins (**Figure 2**). Prokaryotic SMC proteins are large ATPases involved in chromosome organization and segregation (<u>46</u>, <u>47</u>). As with other SMC-like proteins, substantial portions of the SMC-HicA proteins consist of α-helices that fold into coiled-coil structures (**Figure S14A**, **S14B**). The Class 14 HicA domains exhibiting the canonical dsRBD configuration are fused to the C-terminal ends of the SMC domains (**Figure S14B**, **S14C**). Interestingly, some of the SMC-HicA hybrid proteins also have a NERD-domain predicted to have nuclease activity and may function in DNA processing (<u>48</u>).

The function of the HicA domain fused to an SMC homolog remains enigmatic. However, recent discoveries have highlighted the roles of systems like Wadjet and Lamassu (also known as DdmABC in *Vibrio cholerae*) in defending against phage attack and plasmid transformation (<u>1</u>, <u>5</u>) (<u>Robins WP, Mekalanos JJ et al., 2023 bioRxiv</u>). These systems encode SMC homologs that are essential for defense activities. The SMC homologs can be activated by specific signatures in incoming DNA elements. Depending on the system, they either degrade foreign DNA (WadJet) or induce abortive infection (Lamassu) upon detecting invasive elements (<u>5</u>, <u>49</u>) (<u>Robins WP, Mekalanos JJ et al., 2023 bioRxiv</u>). It is thus possible that the ribonuclease activity in Class 14 SMC-HicA proteins might similarly be triggered by foreign DNA or RNA, potentially playing a role in abortive infection or degradation of invasive elements.

### HicA-Encoding Genes are Often Located in Defense Islands

Prokaryotic Defense Islands are regions enriched in antivirus defense systems and mobile genetic elements (MGEs), including prophages, integrative and conjugative elements, transposons, recombinases and IS sequences (<u>2</u>, <u>3</u>). Except for the CRISPR-Cas systems, different classes of defense systems, including TA and restriction-modification systems, exhibit clustering in Defense Islands. To further strengthen the argument that HicA-domains function in phage defense, we engaged in identifying HicA-domains encoded by Defense Islands. The newly developed online tool TADB3.0 facilitates the identification of TA loci encoded by MGEs. As an example, TADB3.0 has cataloged 93 strains of *Shigella flexneri*, among which 40 harbor *hicAB* or *hicBA* loci within MGEs (<u>50</u>). A comparison of Defense Islands from different *Shigella flexneri* strains encoding *hicAB* is shown in **Figure S15A**. Thus, TAs encoding HicA domains are often located within Defense Islands.

### Toxin – Antitoxin Genes Often Cluster

During the manual inspection of DNA sequences encoding *hicA* and *hicB* genes it often became apparent that other TA loci were encoded by neighboring regions. In the archaeon *Methanosarcina barkeri* 3, a Defense Island encodes two *hicBA* loci, two solitary *hicB* genes and one *vapBC* locus (**Figure S15B**). In a *Thermodesulfobacterium hydrogeniphilum* strain, a Defense Island encodes three *hicBA*, three *relBE*, two *vapBC* and a solitary *vapC* gene, in total 8 different TA loci in a region of 7 kilobases of DNA. Furthermore, TA loci often reside in close proximity to transposases or IS elements (<u>50</u>), consistent with their high lateral mobility in Defense Islands.

### Phylogeny of HicA Toxins

The phylogenetic analysis of HicA sequences, spanning Classes 1 to 10, yield insights into the evolutionary relationships among the toxins. Significantly, members of each class tend to form clusters, reflecting their shared structural and functional characteristics, except for Class 4, which bifurcates into two distinct clusters (**Figure 7**). Classes 1 and 2 appear to evolve independently, hinting of separate evolutionary trajectories, despite their similar sizes and the absence of a DNA-binding domain. Similarly, Classes 7 and 8 constitute entirely distinct branches on the tree, consistent with their distinct HicBs. Conversely, the HicAs of Classes 5 and 6, as well as Classes 9 and 10, exhibit a clear pattern of clustering with robust statistical support, suggesting a close genetic relationship between these pairs. However, it is important to note that due to the relatively small size and high sequence variability of HicA domains, the statistical significance of certain branches in the phylogenetic tree may be somewhat limited.

**Figure 7.**
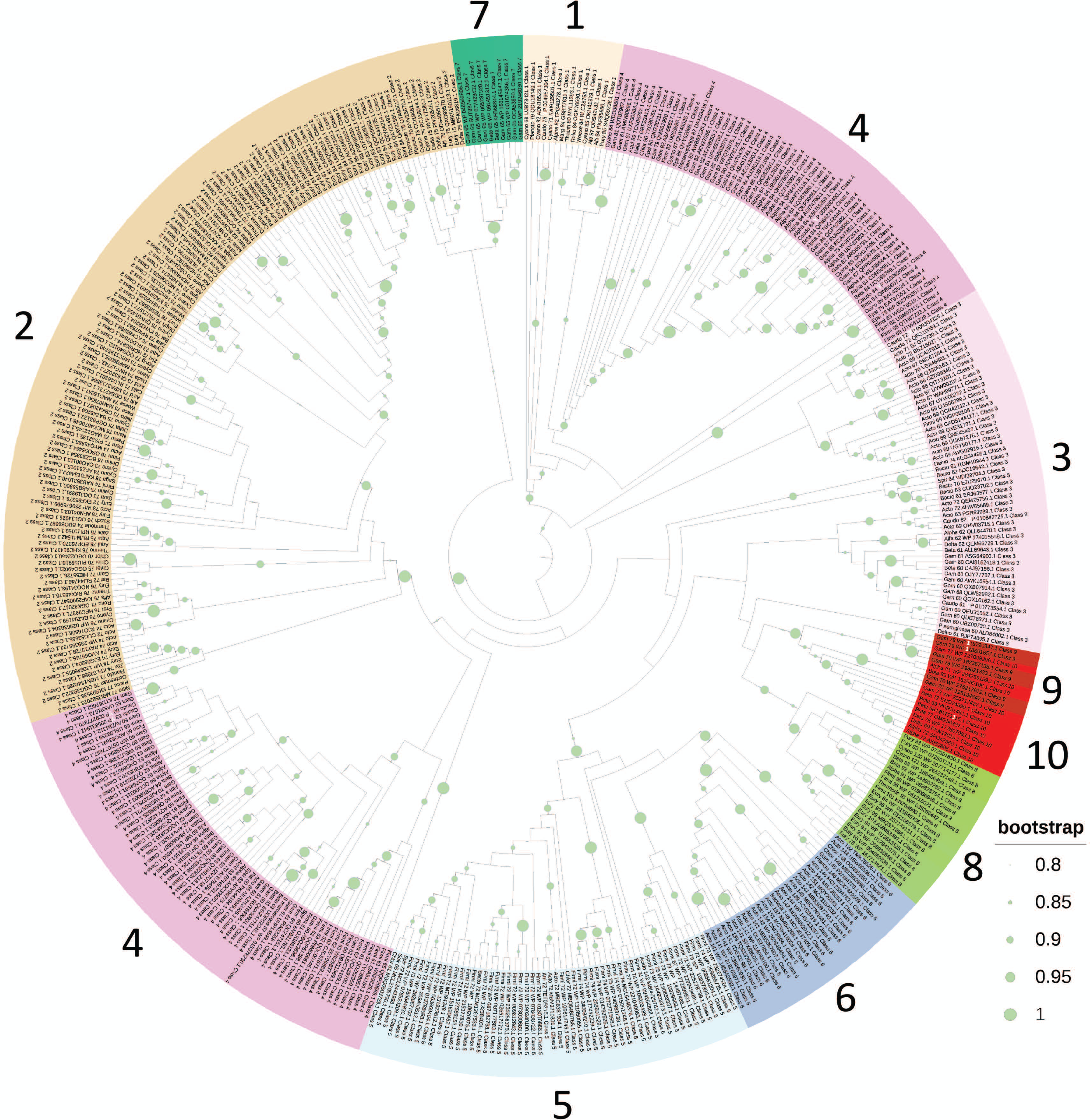
Phylogenetic tree of Class 1 to 10 HicAs. HicA sequences were aligned by Kalign, tree reconstruction performed by the FastTree module of Geneious Prime and visualized by iTOL. Numbers refer to Class 1 to 0. The tree was pruned for outliers.

### Conclusion

The comprehensive analysis of HicA toxin domains has uncovered a diverse array of genetic organizations and functional associations. This study not only identified Class 3 and 4 but also introduced 12 new gene classes encoding HicA toxin domains, significantly expanding our understanding of these systems. Several noteworthy findings and questions have emerged from this investigation: Classes 1 and 2, being the most prevalent, raise the question about the regulation of their gene expression, particularly considering the absence of an identifiable DNA-binding domain in HicB; the reversal in gene order between Classes 3 and 4; the reversal in gene order and transition from HTH/RHH DBD to Phd/YefM DBD in Classes 5 and 6 highlights the variations in DNA-binding modes associated with these gene classes; the identification of the Shared Domain in Class 6 HicA and some RelEs suggests potential functional connections between *hicBA* and *relBE* systems, consistent with a common role in phage-defense; the proximity of Class 9 and 10 HicA-encoding genes to pVip-encoding genes, as well as the occasional association of RelE-encoding genes with pVip genes, also suggests potential cooperation in anti-phage defense; classes 11 to 13 HicA domains are encoded by fused *hicAB* or *hicBA* monocistronic operons, resembling the anti-phage gene CapRelSJ46 and these arrangements are also consistent with a mechanism for curbing phage infections; Class 14, featuring an SMC domain fused to a HicA domain, might also play a role in defense against phages, plasmids, or other mobile genetic elements, akin to the Wadjet and Lamassu systems although alternative functions cannot be ruled out. In conclusion, this study provides substantial support the proposition that genes encoding HicA domains play pivotal roles in defense against phages, plasmids, and other mobile genetic elements.

## Materials and Methods

### Data Sampling

Sequences of experimentally validated HicA toxins were used as seeds in BLASTP searches at NCBI (https://blast.ncbi.nlm.nih.gov/), using different bacterial phyla as search spaces (**Table S1**). HMMSEARCH at <u>ebi.ac.uk</u> (<u>51</u>) was used to expand poorly populated Clades. In **Table S1**, kept HicA sequences are less than 95% identical to any other kept HicA sequence.

**Toxin – Antitoxin Gene Organization and Gene Neighborhood Analysis** were accomplished using webFlaGs (<u>52</u>) or TADB3.0 (<u>50</u>). The gene organizations shown in **Figure 2** were then confirmed by manual inspection of the DNA regions encoding the HicAs of **Table S1**.

### Sequence Alignments and Phylogenetic Trees

Sequence alignments were generated by Clustal Omega (<u>53</u>) or Kalign (<u>54</u>) at www.ebi.ac.uk and imported into Jalview (<u>55</u>). Protein sequence alignments in Jalview 2.11.0 were exported as vector files (EPS or SVG formats), converted to jpg files and imported into Adobe Illustrator, annotated and saved in PDF format for publication. Phylogenetic trees were visualized using iTOL (<u>56</u>). Reconstruction of phylogenetic trees was accomplished using FastTree (<u>57</u>) via the CIPRES module in Genious Prime that uses the Maximum Likelihood approach and Ultrafast bootstrapping.

### Protein Structure Prediction, Protein Similarity Searches and Protein Structure Visualization

Protein secondary structures were predicted from sequence alignments using the link to JPred (<u>58</u>) in JalView. Protein tertiary structures were modelled using AlphaFold2 (<u>59</u>) via the ColabFold v1.5.2-patch (<u>60</u>) or the AlphaFold patch of ChimeraX. Mutimeric structures were modelled by MultiFOLD (<u>61</u>) or AlphaFold2 and validated by ModFOLDdock (<u>62</u>). Structure similarity searches was done using Phyre2 (<u>63</u>) or FoldSeek (<u>64</u>). Structures were visualized and annotated using ChimeraX (<u>Pettersen *et al.*, 2021</u>). HTH motifs were identified by EMBOSS (<u>65</u>).

### Categorization of HicB Antitoxins Encoded by Class 1 *hicAB* Operons

Class 1 *hicB* genes encode HicB lacking a DBD. These *hicB* genes could encode either a functional protein or a pseudogene that had arisen by mutation, e.g. a stop-codon mutation that truncated the *hicB* gene. To analyze if this was the case, each individual Class 1 HicB protein sequence was used as a query in BLASTP searches at NCBI (https://blast.ncbi.nlm.nih.gov/). Some HicBs were almost identical to a HicB with a DBD; operons of this type were discarded. However, most HicB antitoxins exhibited a BLASTP search pattern very similar to other HicBs, revealing homologs of similar sizes but with more distantly related protein sequences, indicating that these proteins were actively evolving even though they lacked a DBD. These proteins were included in the Class 1 *hicBA* operons. Furthermore, the relative uniform size distribution of Class 1 HicBs (60 to 87 aa; **Table S1**) is consistent with the notion that the antitoxins are not products of randomly mutated *hicB* genes.

## Supplemental Material

### Legends of Supplementary Figures

**Figure S1.**
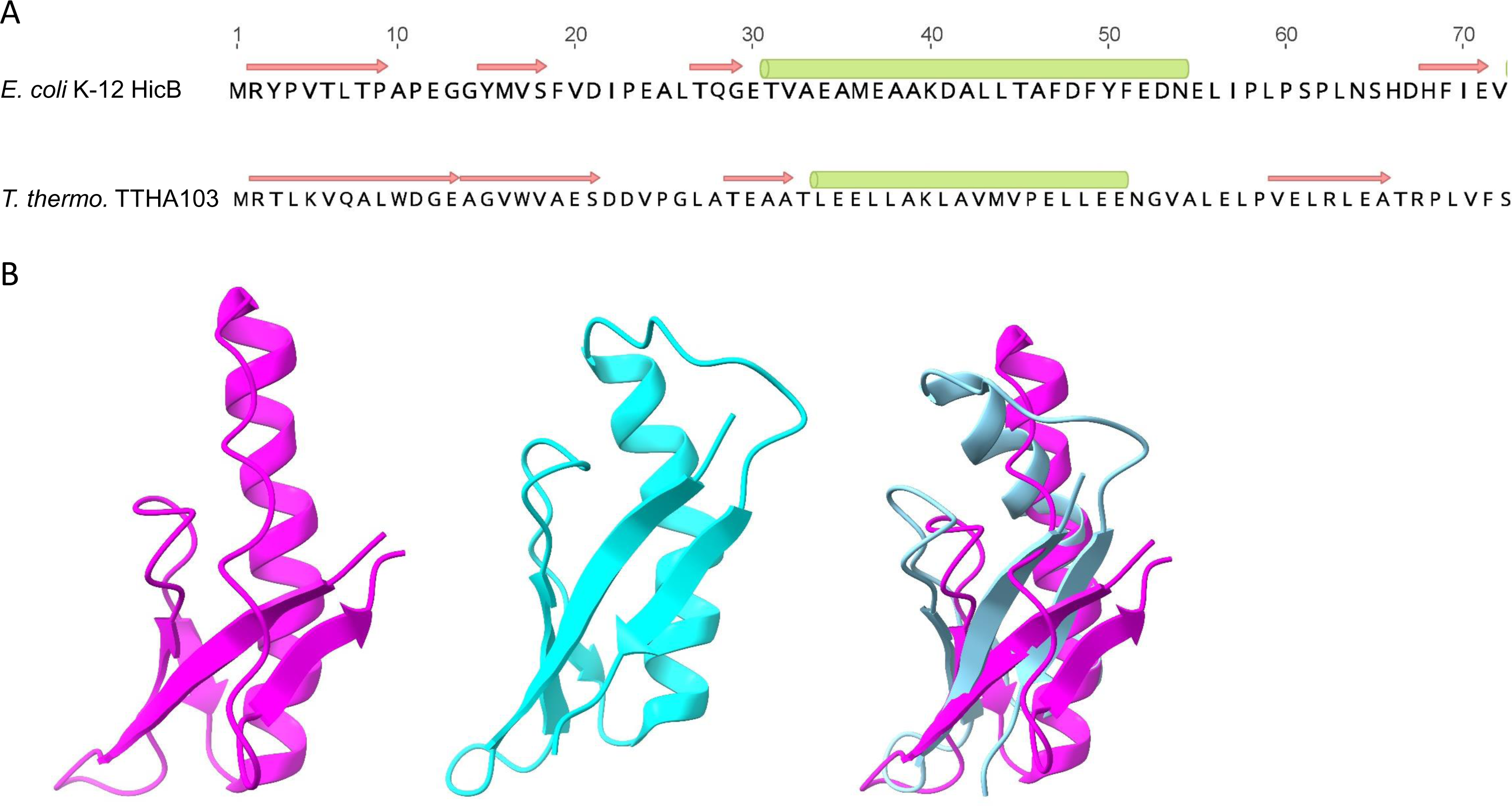
HicB of *E. coli* K-12 exhibits a partial RNase H fold. (**A**) Comparison of experimental secondary structures of HicB of *E. coli* K-12 (P67697.2, upper) and *T. thermophilus* HB8 (BAD71735.1, lower). (**B**) Tertiary structures of the N-terminal partial RNase H-like domain of HicB of *E. coli* K-12 (6HPB.pdb, left), TTHA1013 of *T. thermophilus* HB8 middle and their superimposition (right) with an RMSD between 18 pruned atom pairs of 1.4Å. These structures were obtain experimentally (<u>1</u>, <u>2</u>).

**Figure S2.**
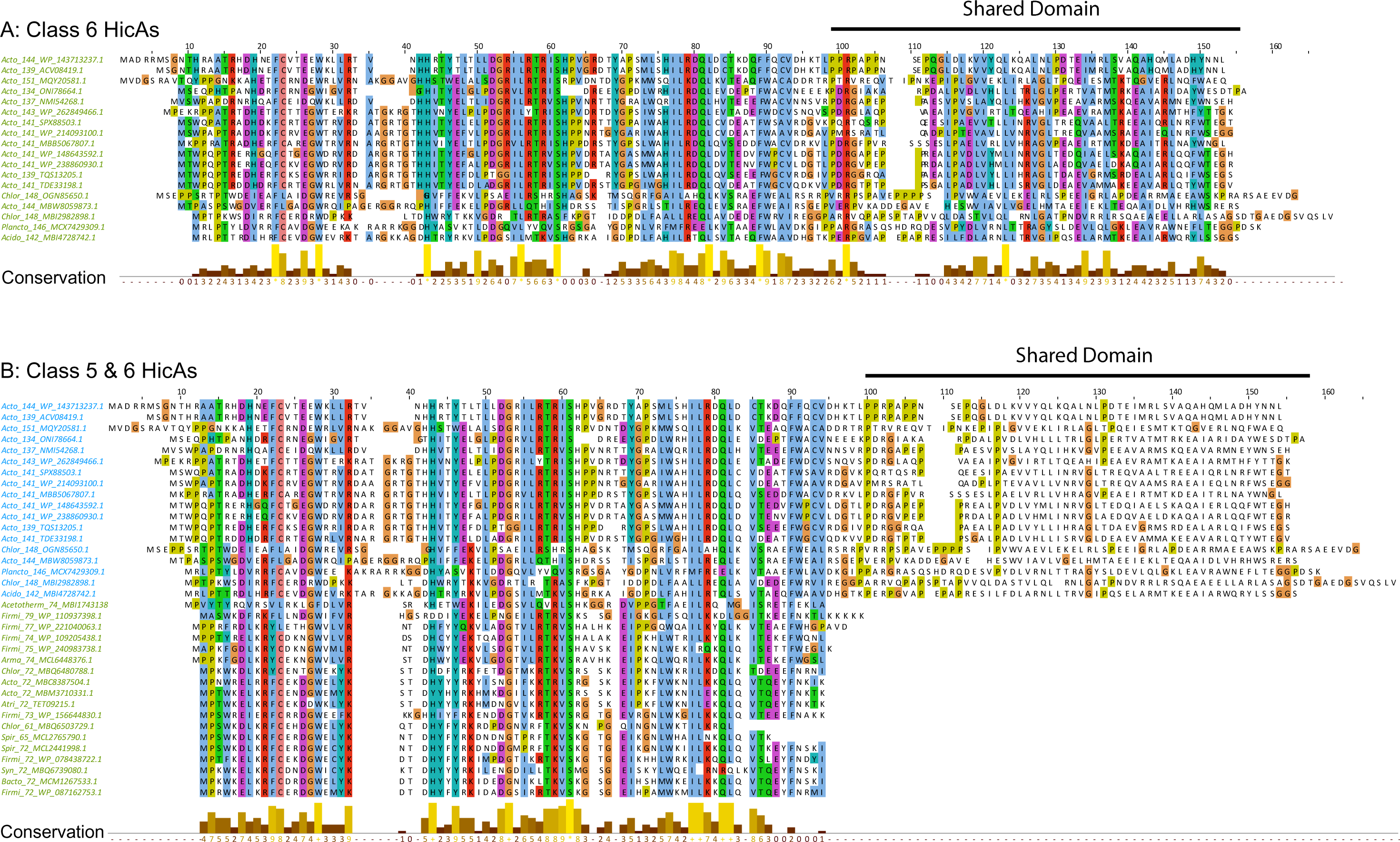
Sequence alignments of Class 5 and 6 HicA. (**A**) Alignment of Class 6 HicAs. The Shared Domain is indicated with a bar. (**B**) Alignment of Class 5 and 6 HicAs. Class 5 HicAs are typical small, mono-domain proteins with a dsRBD (**Table S1**) and while Class 6 HicAs are two domain proteins consisting of a typical N-terminal dsRBD and a second domain here called the Shared Domain of 50 to 55 amino acids. Conserved proline residues (green) may serve as domain-breakers between the two domains.

**Figure S3.**
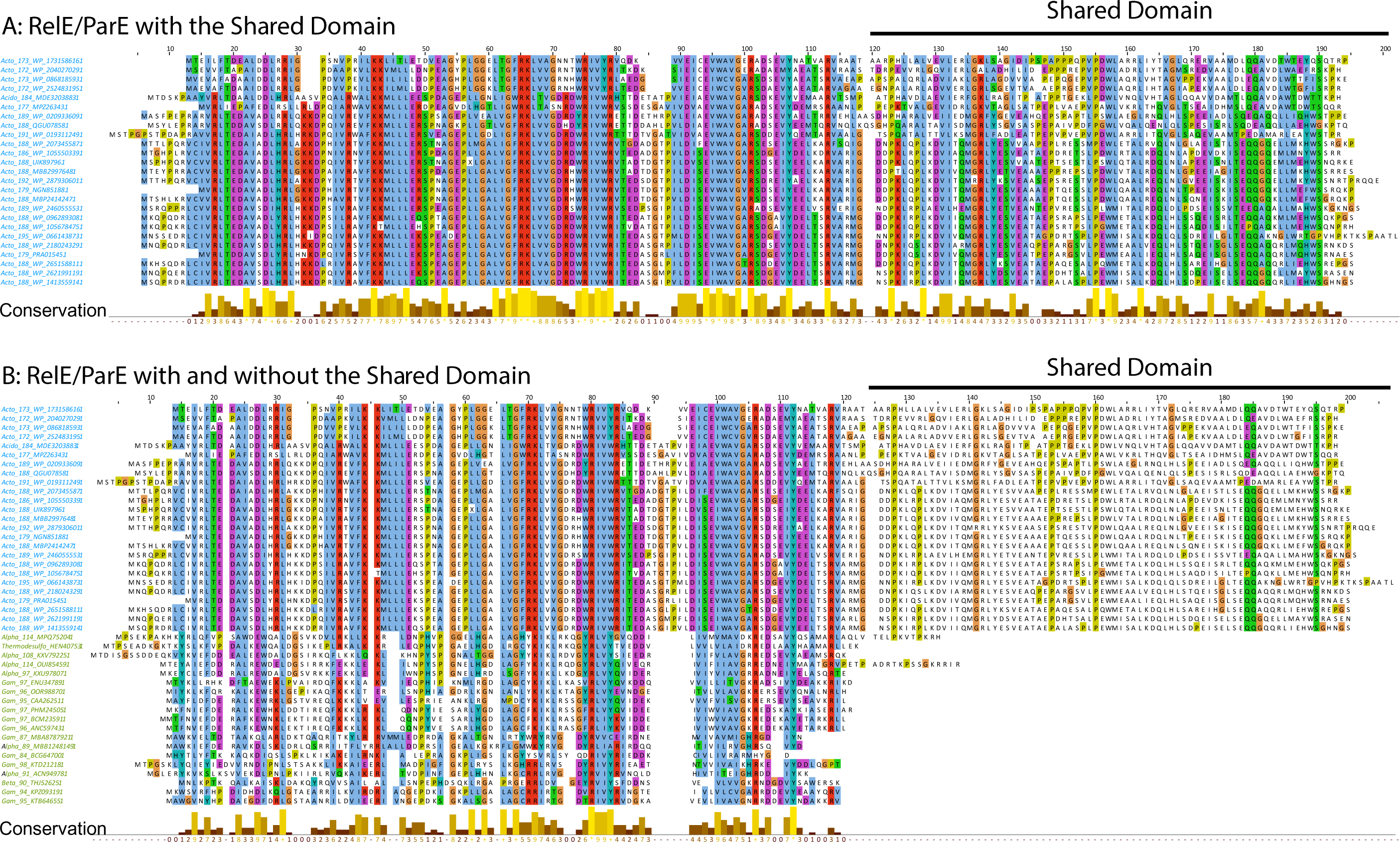
Sequence alignments of RelE/ParE toxins with and without the Shared Domain. (**A**) Alignment of RelE/ParE superfamily of toxins with the typical N-terminal RNase I domain and a Shared C-terminal Domain (70 to75 aa). (**B**) Alignment of typical mono-domain RelE/ParE toxins and RelE/ParE toxins with a Shared C-terminal Domain.

**Figure S4.**
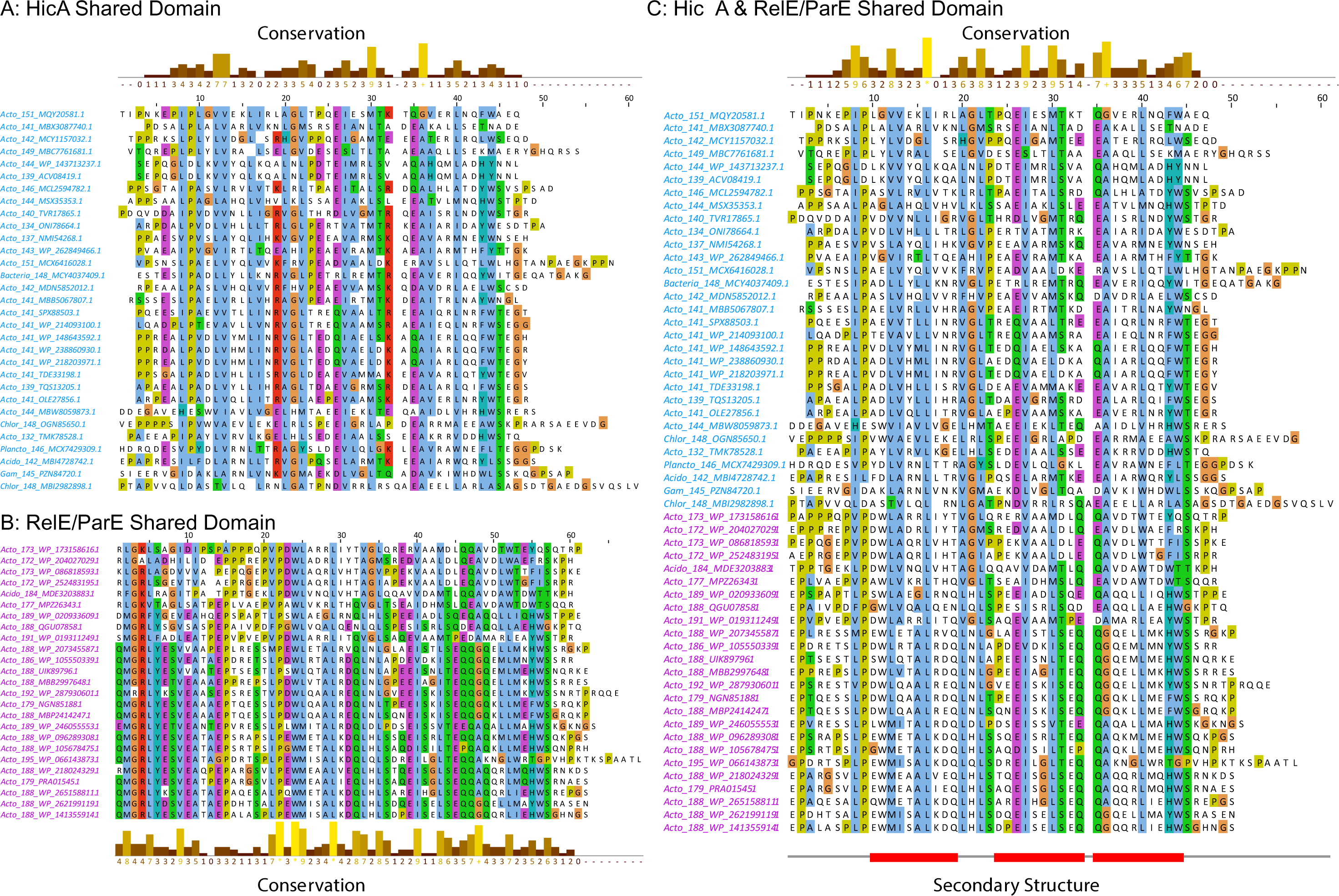
Sequence alignments of the Shared Domains of Class 6 HicAs and RelE/ParE. (**A**) Alignment of the Shared Domains of Class 6 HicAs. (**B**) Alignment of Shared domains of RelE/ParE family toxins. (**C**) Alignment of the two classes of Shared Domains.

**Figure S5.**
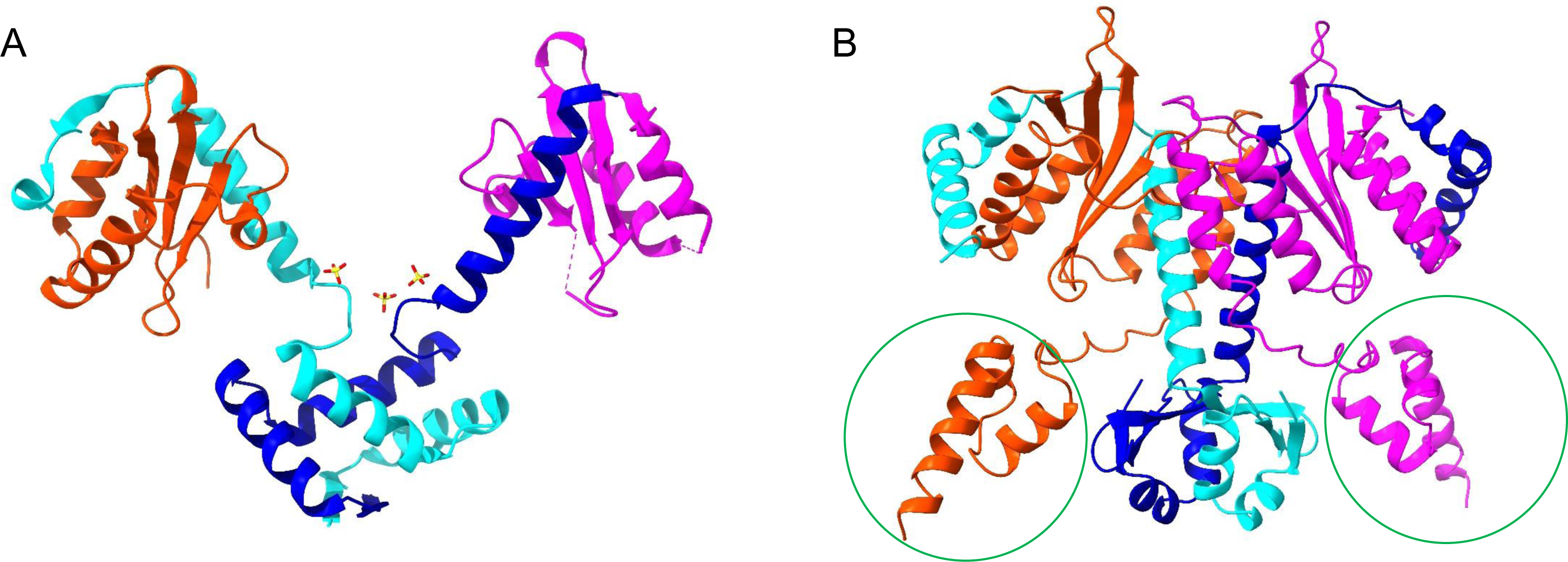
Comparison of Quaternary Structure Models of RelB_2_RelE_2_ Complexes. (**A**) Experimental RelB_2_E_2_ tetramer model of canonical RelE (CAA26251.1) and RelB (CAA26250.1) of *E. coli* K-12 (<u>3</u>). (**B**) RelB_2_E_2_ tetramer model in which RelE has a Shared Domain (WP_141355914.1) and RelB (WP_174787664.1) of *Glutamicibacter nicotianae* (an Actinomycete) generated by MultiFOLD (pLLDT = 0.67). The Shared Domains are marked with green circles. Note that RelB in (A) dimerizes via a HTH domain while RelB in (B) dimerizes via a Phd/YefM Domain.

**Figure S6.**
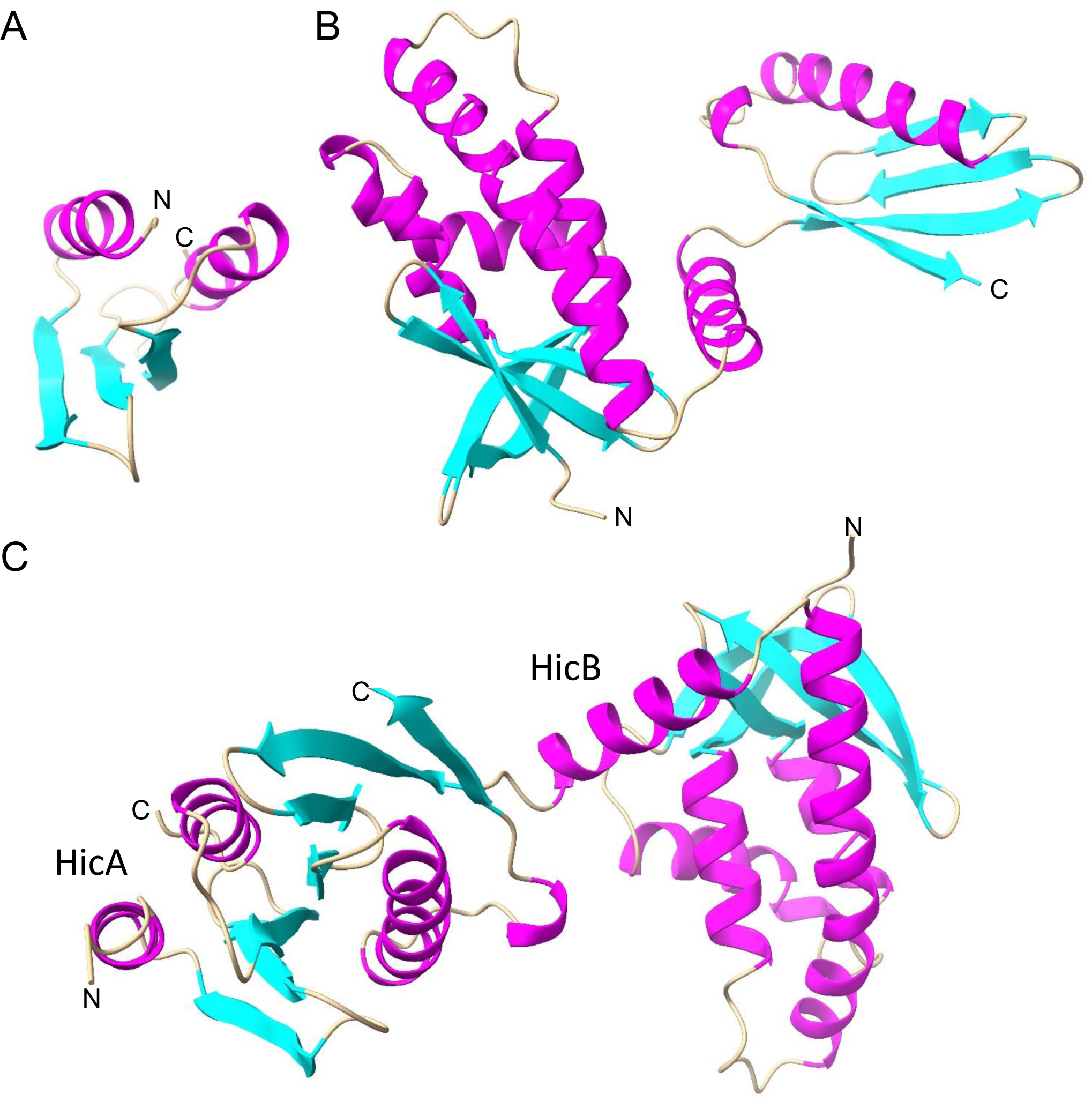
Tertiary and Quaternary Structure Models of Class 7 HicA, HicB and HicAB. (**A**) Predicted model of Class 7 HicA (WP_001674386.1; 65 aa) revealing the canonical dsRBD α-β-β-β-α fold of HicA. (**B**) Predicted Model of Class 7 HicB (WP_000065196.1; 217 aa) revealing its two-domain structure. (**C**) Dimer structure of Class 7 HicA and HicB modelled by MultiFold. Dimer plDDT: 0.918; Dimer pTM: 0.750; Assemby quality: 0.9098; Interface quality: 0.9058.

**Figure S7.**
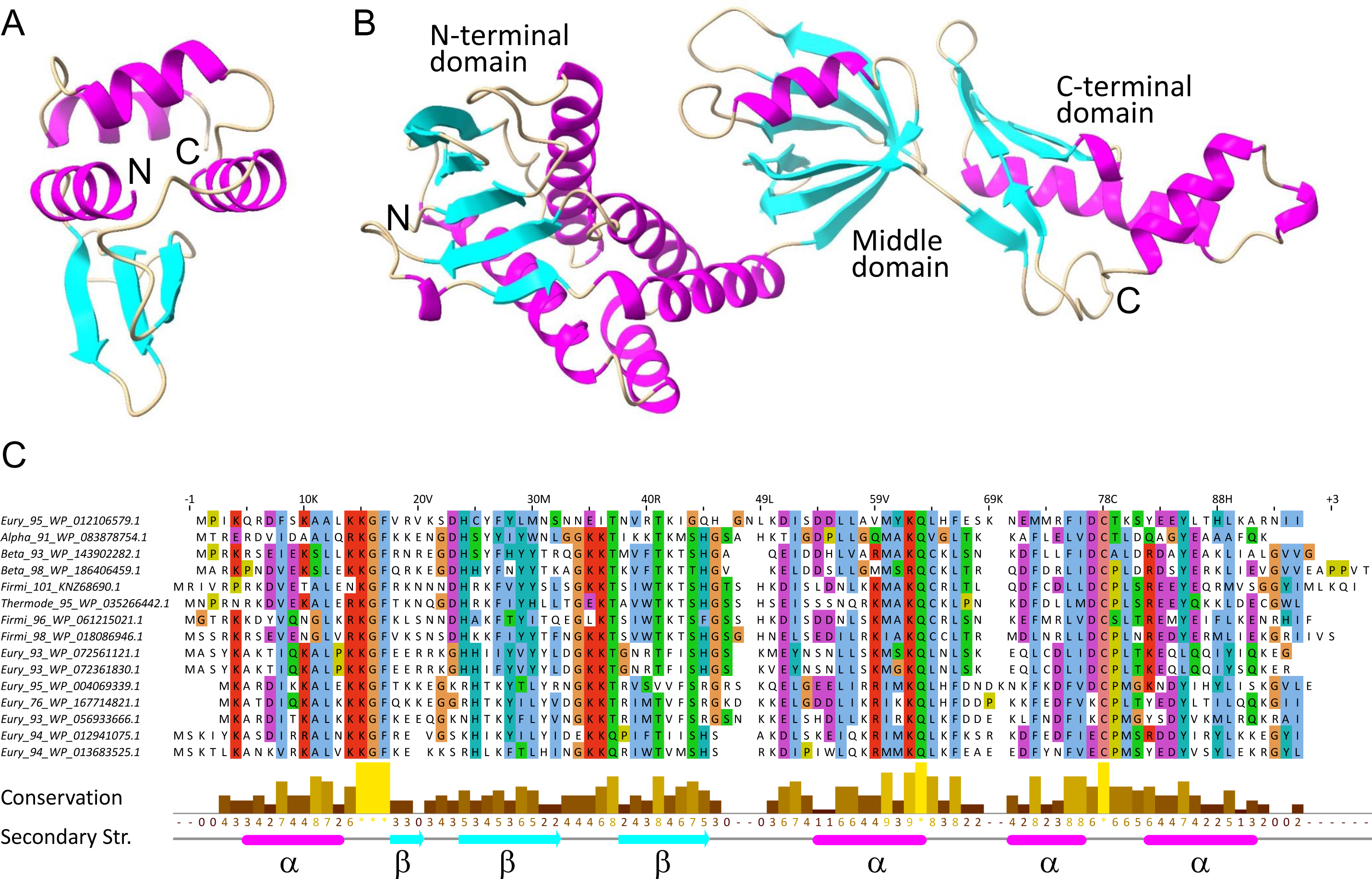
Secondary and Tertiary Structures of Class 8 HicA and HicB. (**A**) Tertiary structure of Class 8 HicA (GenBank ID: WP_004069339.1; 95 aa) modelled by AF2. (**B**) Tertiary structure of Class 8 HicB (GenBank ID: WP_004069337.1; 342 aa) modelled by AF2. See **Table S1** for further details. (**C**) Sequence alignment and secondary structure prediction of Class 8 HicA sequences. The prediction was generated by analysis of an alignment of Class 8 HicA sequences in the JPRED module of Jalview. α-helices are colored red and β-sheets green.

**Figure S8.**
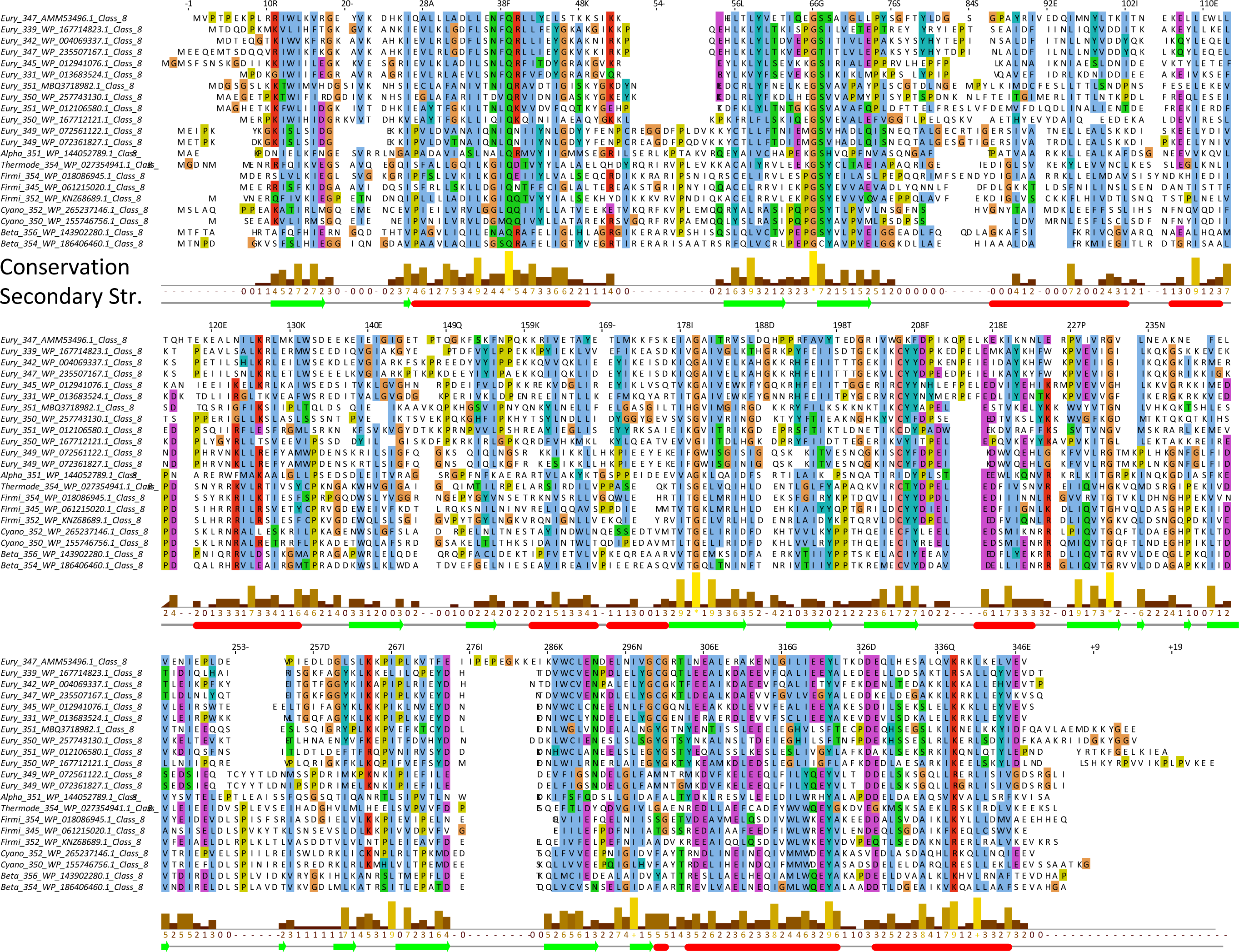
Sequence Alignments and Secondary Structures of Class 8 HicAs and HicBs. Sequence alignment and secondary structure prediction of Class 8 HicB sequences. The prediction was generated by analysis of an alignment of Class 8 HicA sequences in the JPRED module of Jalview. α-helices are colored red and β-sheets green.

**Figure S9.**
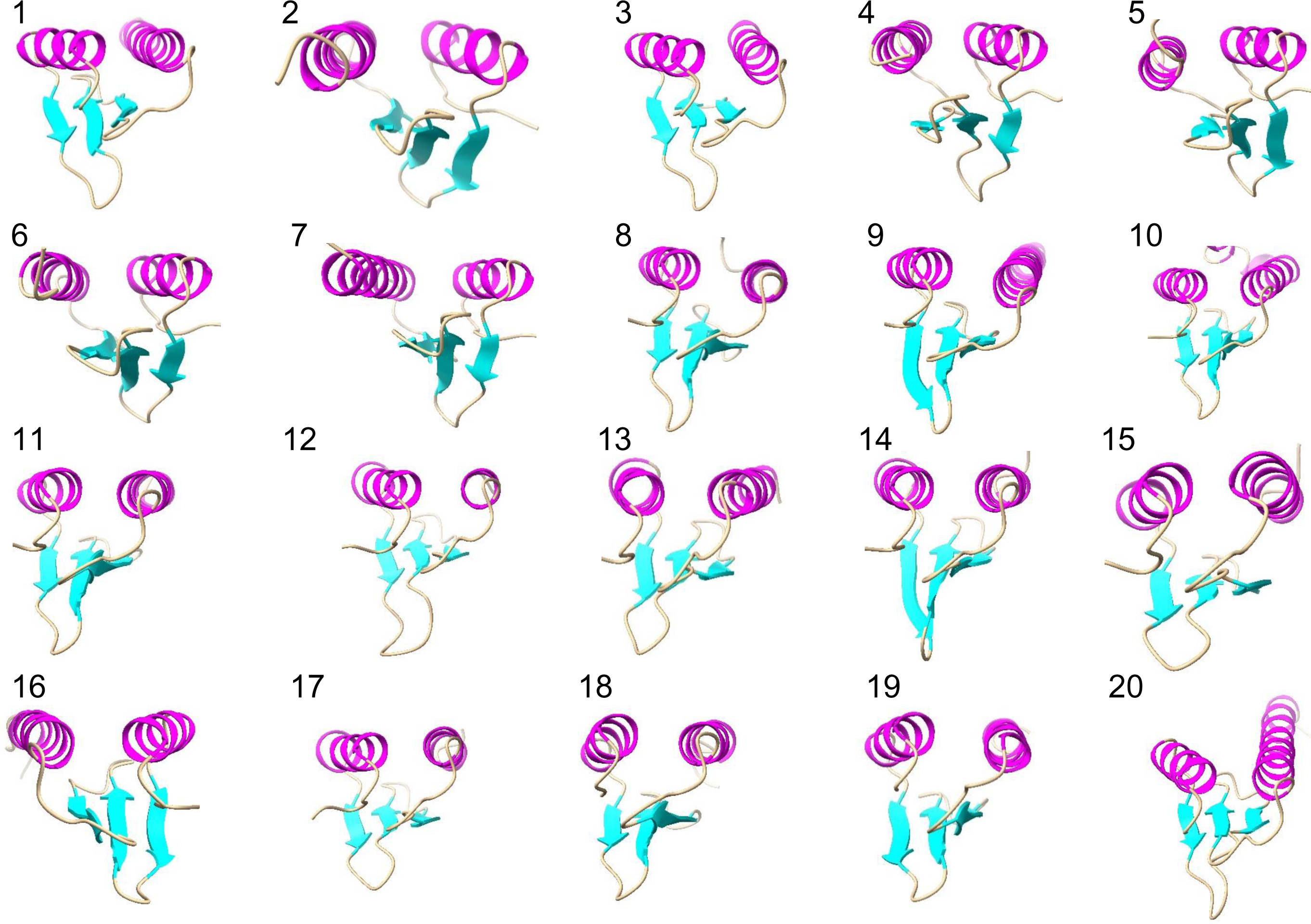
Tertiary Structure Models of Class 10 HicAs. The tertiary structures of 21 Class 10 HicAs were modelled be AlphaFold2. They all have a pLLDT score > 90. Numbers refer to the listing in **Table S1** (Sheet 19).

**Figure S10.**
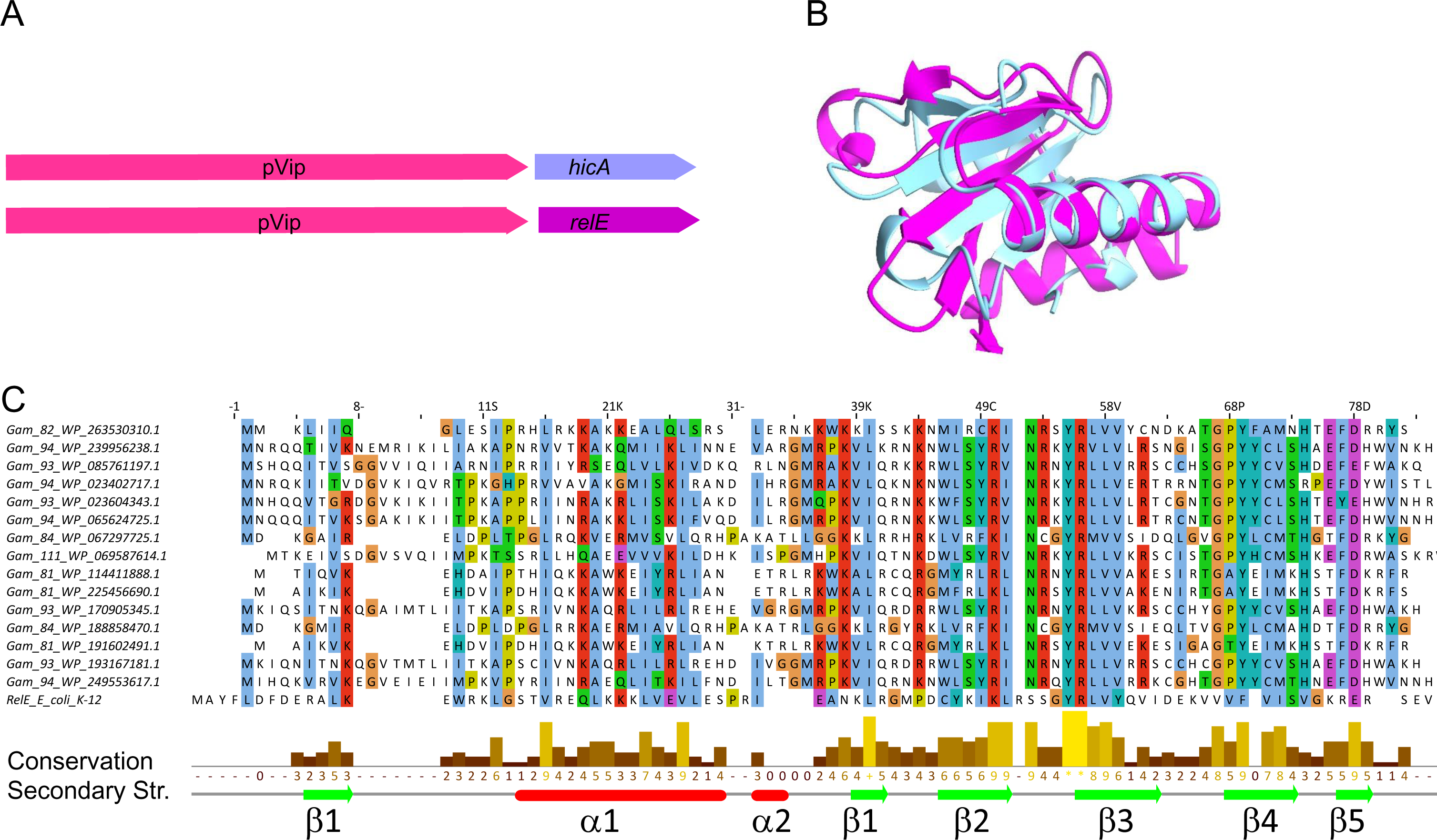
RelE-Encoding Genes Adjacent to pVip Genes. (**A**) Comparison of the organization of genes encoding pVip and HicA, and pVip and RelE. (**B**) Superposition of RelE from a pVip-encoding operon (WP_263530310.1; cyan) and RelE-homolog YoeB of *E. coli* K-12 determined experimentally (<u>4</u>) (2A6S; magenta). The RMSD between 19 pruned atom pairs is 0.835 Å. (**C**) Sequence alignment and secondary structure prediction of 15 RelE-homologous RNases encoded by genes juxtaposed to pVip-encoding genes and RelE of *E. coli* K-12 (bottom).

**Figure S11.**
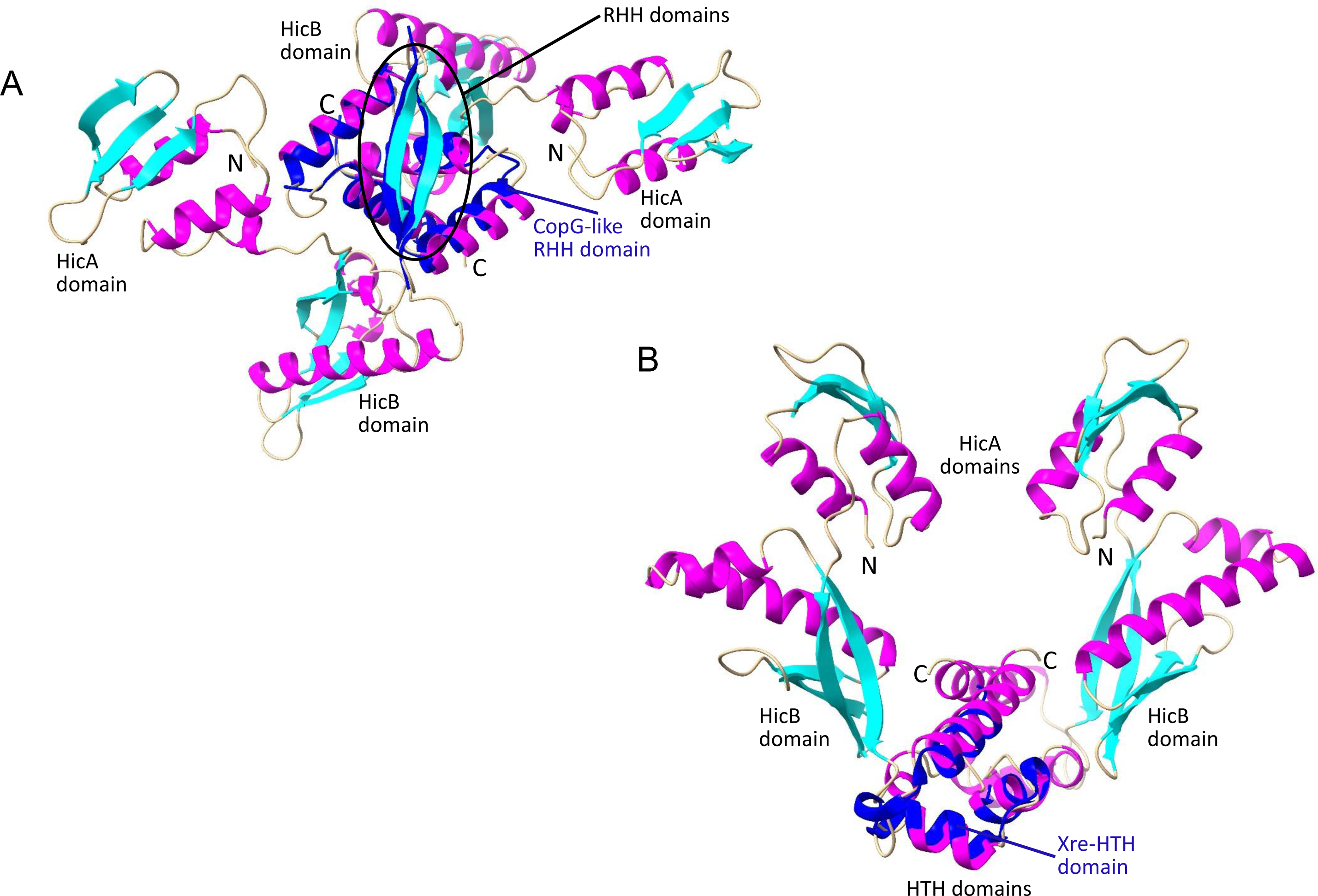
Class 11 Fused HicAB Dimer Model Structures. (**A**) Predicted dimer model of Class 11 fused HicAB (KPW96986.1). The monomers dimerizes via their RHH domains that are superimposed on a plasmid-encoded CopG dimer shown in blue (<u>5</u>) (RMSD between 25 pruned atom pairs is 0.753Å). (**B**) Predicted dimer model of fused Class 11 HicBA (WP_243550782.1). The monomers dimerizes via their HTH domains that are superimposed on the HTH domain of HipB of *E. coli* K-12 (<u>6</u>) (RMSD between 35 pruned atom pairs is 1.226Å).

**Figure S12.**
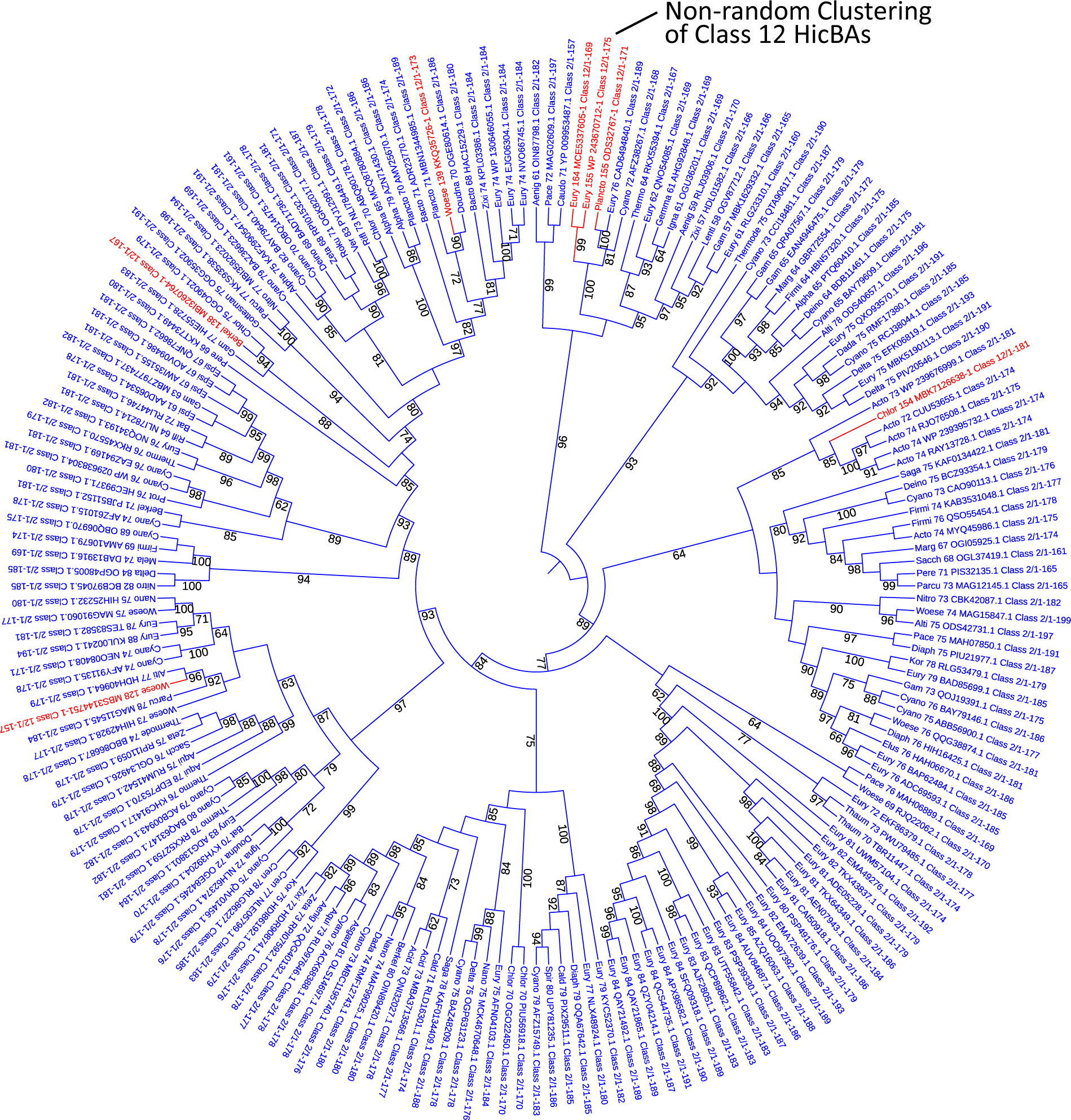
Phylogenetic Tree of 185 Class 2 HicBAs generated by fusion of the HicB and HicA proteins and 7 Class 12 naturally fused HicBAs. The HicBA sequences were aligned with Clustal Omega and the Tree generated by the FastTree module of Geneious Prime using Ultrafast bootstrapping.

**Figure S13.**
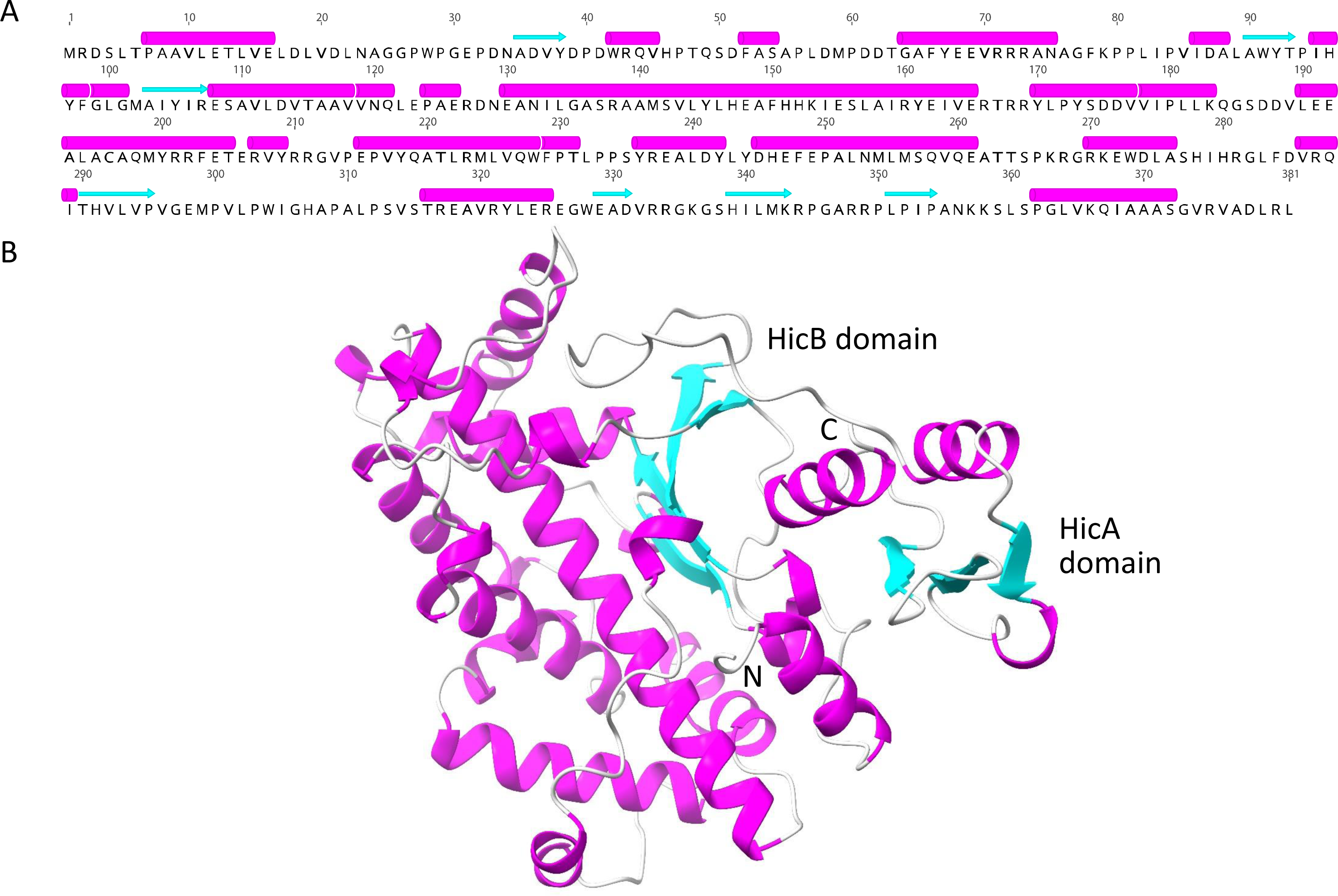
Secondary and Tertiary Structures of a Fused Class 13 HicBA Toxin – Antitoxin. (**A**) Secondary structure of Class 13 HicBA (WP_067334119.1). Helices are in magenta, sheets in cyan and coil in light grey. (**B**) Tertiary structure of Class 13 HicBA (WP_067334119.1).

**Figure S14.**
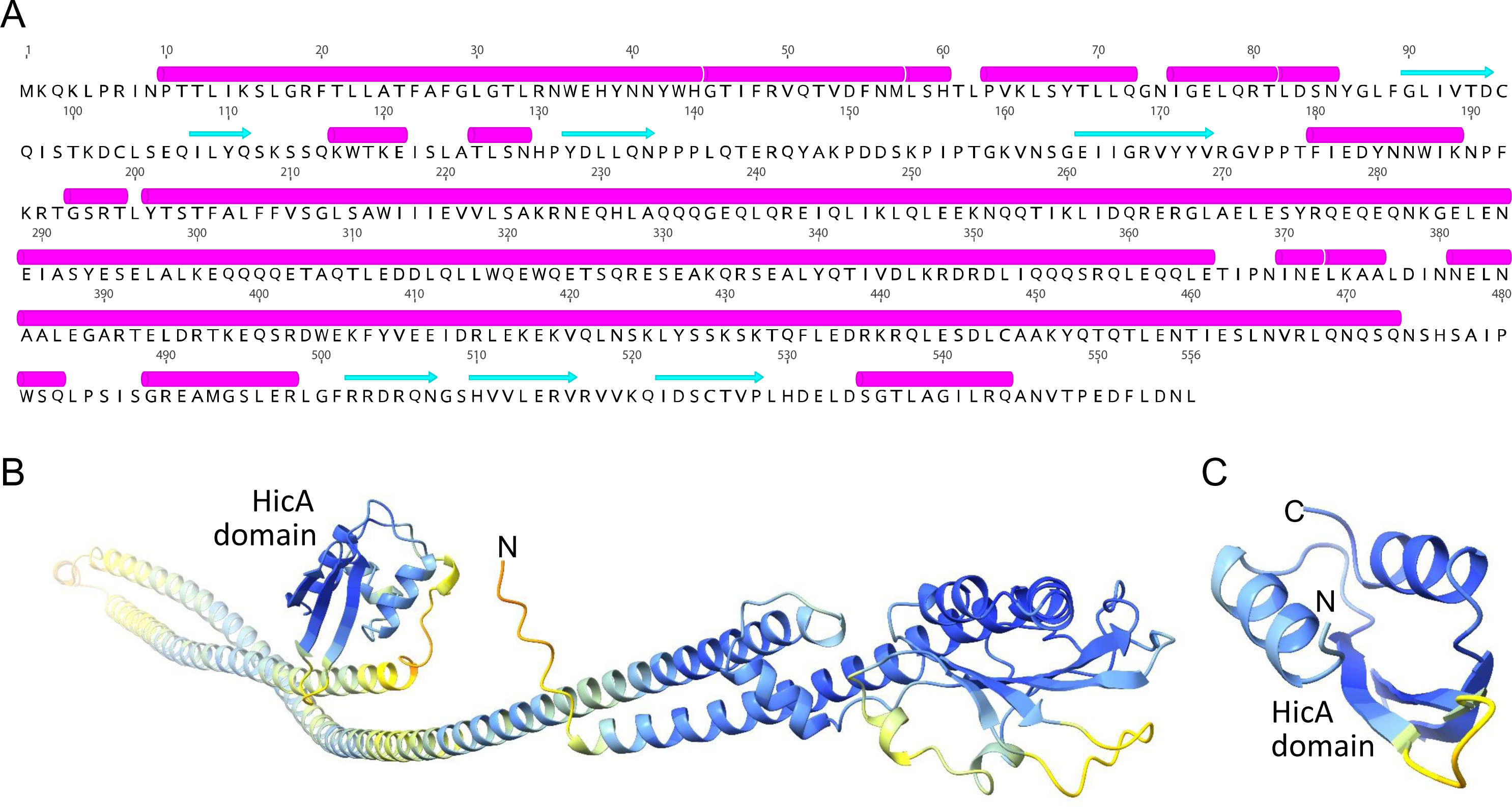
Predicted Secondary and Tertiary Structures of a Fused SMC-HipA Protein. (**A**) Primary and secondary structures of a Class 14 SMC-HicA two-domain protein (WP_223300692.1). Helices are in magenta, sheets in cyan and coil in light grey. (**B**) Tertiary structure of Class 14 SMC-HicA protein (WP_223300692.1). The coloring palette was from AlphaFold2 where blue is high and yellow low quality of the prediction. In particular, loops and parts of the coiled-coil domains were predicted with low quality whereas the C-terminal HicA domain and a globular domain were predicted with high quality. (**C**) Enlargement of the C-terminal HicA-domain of the structure shown in (B).

**Figure S15.**
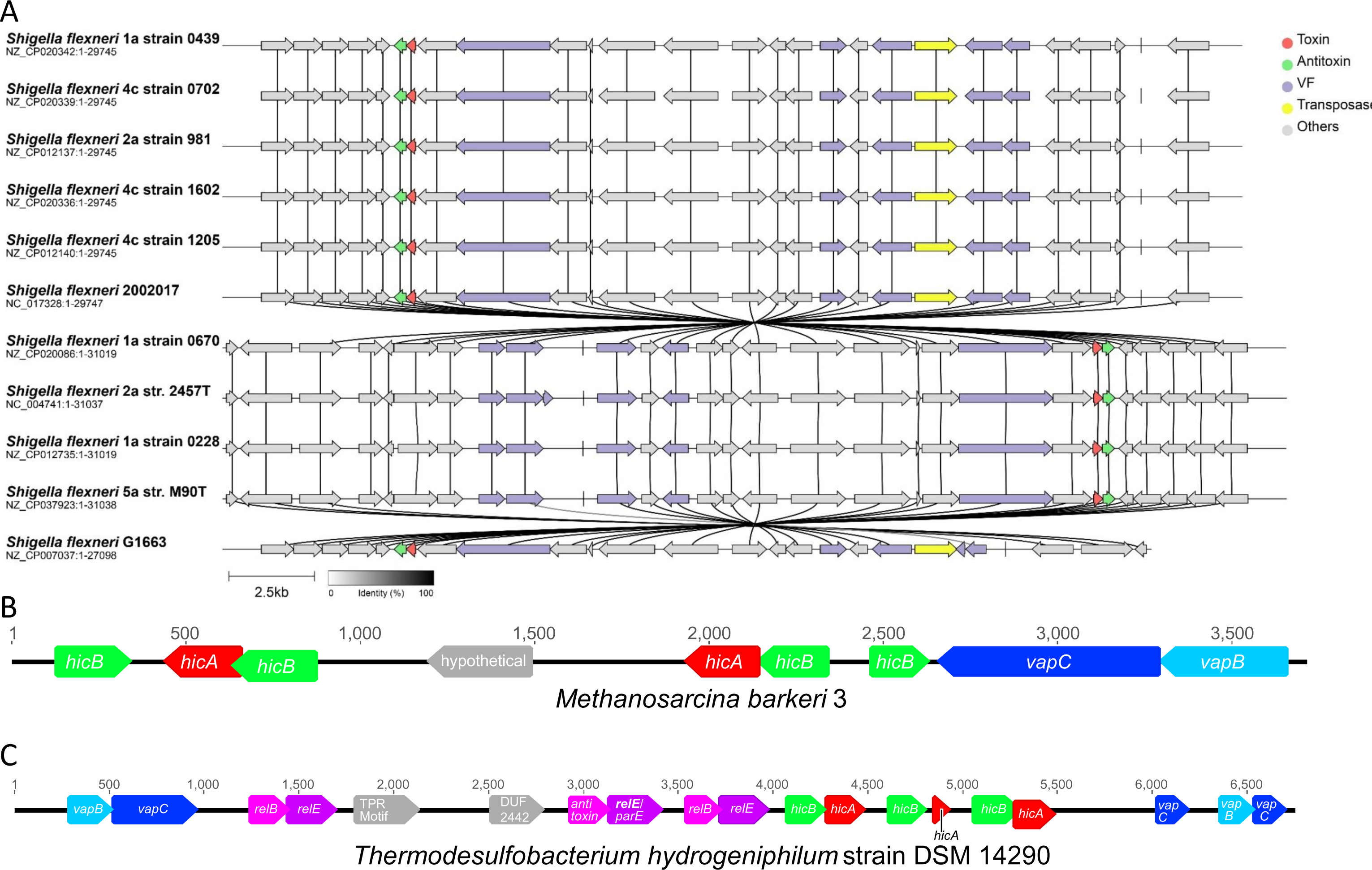
Defense Islands Encoding *hicAB* and *hicBA* Genes. (**A**) Alignment of DNA sequences encoding Defense Islands of 11 *Shigella flexneri* strains encoding *hicAB* loci. The *hicA* and *hicB* genes are colored red and green, respectively, transposase genes are yellow and virulence factors magenta. The data were retrieved from TADB 3.0 and the Figure generated by Clinker & Clustermap (<u>7</u>) embedded in TADB3.0. (<u>8</u>). (**B**) Defense Island of archaeon *Methanosarcina barkeri* 3 (NZ_CP009517.1) encoding two *hicBA* green, red), two solitary *hicB* genes (green) and a *vapBC* locus (pale blue, blue). (**C**) Defense Island of *Thermodesulfobacterium hydrogeniphilum* strain DSM 14290 (NZ_JQKW01000008) encoding three *hicBA* (green, red), three *relBE* (magenta, pink), two *vapBC* (pale blue, blue) and one *vapC* gene (blue). TPR: gene encoding a tetratricopeptide repeat protein. Genes unrelated to TA modules are shown in grey. DUF: domain of unknown function. DUF2442 is often found in antitoxins (<u>9</u>).

### Supplementary Table

**Table S1:** Primary information of members of the 14 Classes of HicA-domain encoding TA modules identified in this work.

**Sheets 1 to 14:** Information of HicA and HicB of the 14 classes. Columns A to J yield the following information: (**A**) GenBank ID of HicA-domain protein; (**B**) Organism; (**C**) Label yielding phylum, gene length in codons, GenBank ID and Class of HicA-domain protein. These labels were used in sequence alignments and in the phylogenetic tree (**Figure 7**); (**D**) HicA sequence; (**E**) Distance between *hicA* gene and its cognate antitoxin-encoding gene in nucleotides (a minus indicates gene overlap); (**F**) Genetic order (GO) of each toxin – antitoxin module; (**G**) HicA Class; (**H**) Label yielding HicB antitoxin phylum, gene length and GenBank ID and Class; (**I**) HicB Antitoxin or pVIPs sequence; (**J**) Domains in HicB and comments to each individual entry in the Table. Full names of phyla are given in **Sheet 17**.

**Sheet 15:** *relBE* / *parDE* modules where the toxins contain the C-terminal Shared Domain also present in Class 6 HicA.

**Sheet 16:** Information of anti-phage modules encoding pVip and RelE toxin.

**Sheet 17:** List of Phyla abbreviations used in Sheets 1 to 16.

